# Balance between direct and indirect pathways of the nucleus accumbens controls social behavior in mice

**DOI:** 10.1101/2022.11.28.518147

**Authors:** J. Le Merrer, B. Detraux, J. Gandía, A. De Groote, M. Fonteneau, A. de Kerchove d’Exaerde, J.A.J. Becker

**Affiliations:** Physiologie de la Reproduction et des Comportements, INRA UMR-0085, CNRS UMR-7247, Université de Tours, Inserm, 37380 Nouzilly, France; Inserm UMR-1253 iBrain, Université de Tours, CNRS, Faculté des Sciences et Techniques, Parc de Grandmont, F-37200 Tours, France; Laboratory of Neurophysiology, ULB Neuroscience Institute, Université Libre de Bruxelles (ULB), 1070 Brussels, Belgium; WELBIO, 1300 Wavre, Belgium

## Abstract

**Background:** Deficient social interactions are a hallmark of major neuropsychiatric disorders, and cumulating evidence point to altered social reward and motivation as key underlying mechanisms in these pathologies. In the present study, we aimed at assessing the role of the two striatal projecting neuronal (SPN) populations bearing either D1R or D2R dopamine receptors (D1R- and D2R-SPNs), in modulating social behavior and other behaviors often altered in neuropsychiatric disorders.

**Methods:** We selectively ablated D1R- and D2R-SPNs using an inducible diphtheria toxin receptor (DTR)-mediated cell targeting strategy and assessed social behavior as well as repetitive/perseverative behavior, motor function and anxiety levels. We tested the effects of optogenetic stimulation of D2R-SPNs in the *Nucleus Accumbens* (NAc) and pharmacological compounds repressing D2R-SPN.

**Results:** Targeted deletion of D1R-SPNs in the NAc blunted social behavior in mice, facilitated skill motor learning and increased anxiety levels. These behaviors were normalized by pharmacological inhibition of D2R-SPN, which also repressed transcription in the efferent nucleus, the ventral pallidum (VP). In contrast, ablation of D1R-SPNs in the dorsal striatum had no impact on social behavior, impaired motor skill learning, and decreased anxiety levels. Deletion of D2R-SPNs in the NAc also produced motor stereotypies but facilitated social behavior and impaired skill motor learning. We mimicked excessive D2R-SPN activity by optically stimulating D2R-SPNs in the NAc and evidenced a severe deficit in social interaction that was prevented by D2R-SPN pharmacological inhibition.

**Conclusions:** Repressing D2R-SPN activity may represent a promising therapeutic strategy to relieve social deficit in neuropsychiatric disorders.

## INTRODUCTION

When conditions are favorable, interactions with conspecifics in social animals are pleasurable (1–3), which in turn fuels social motivation. Compromised social interactions are a hallmark of major neuropsychiatric disorders, such as autism spectrum disorder (ASD), schizophrenia and depression, and cumulating evidence point to altered social reward and motivation as key underlying mechanisms in these pathologies (4–7). However, the neuronal and molecular underpinnings of social abilities remain scarcely known. Better delineating these substrates is necessary to develop innovative pharmacotherapeutic strategies able to restore social function when pathologically impaired.

The identification of brain regions and specific networks underlying social processing in social animals has emerged and consolidated over the past 30 years (8–10). Interestingly, this social brain widely overlaps with the brain reward circuit (11–13). Within this circuit, the NAc works as a hub structure, integrating multiple and complex inputs from cortical, allocortical, thalamic, midbrain and brainstem regions (14–16) and, as such, plays a unique role in reward processing and approach (15, 17, 18). Of interest, the view of the NAc as a key substrate for modulating social behavior is spreading in clinical and preclinical literature. Imaging studies have revealed NAc activation in response to various social stimuli in humans (19, 20), activation that was found blunted in patients with ASD, depression or schizophrenia (21–23). Consistently, anomalies in the volume and/or connectivity of the NAc have been evidenced in patients with autism, schizophrenia or depression (24–28) as well as in preclinical models of these pathologies (29–34). Dopamine (DA) (35), serotonin (36), opioid (37–39) and oxytocin (40, 41) receptor activation in the NAc was shown to stimulate/drive social behavior in rodents. Thus, the NAc appears as a key brain structure for the control of social abilities under physiological and pathological conditions. How the two main neuronal populations in the NAc, D1R- and D2R-SPNs, reciprocally contribute to modulating social behaviors, however, remains barely understood.

D1R- and D2R-SPNs differ in their transcriptional and projection patterns. Throughout the striatum, D1R-expressing SPNs also express *Pdyn* (coding for prodynorphin) and *Tac1* (for Substance P) whereas D2R expression is associated with *Penk* (for proenkephalin), *Adora2a* (for adenosine 2a receptor) and *Gpr6* (for orphan receptor GPR6) (42–45) expression. D1R and D2R SPNs project to distinct brain areas. In the dorsal striatum (DS), D1R SPNs project directly to the output nuclei of the basal ganglia, the substantia nigra pars reticulata (SNr) and internal globus pallidus (GPi, or entopeduncular nucleus in rodents), making the direct pathway; D2R-SPNs project to the external globus pallidus (GPe) that projects to the subthalamic nucleus (STN), which in turn innervates the output nuclei SNr and GPi/EP, forming the indirect pathway (46). In the NAc, D1R-SPNs project to the VTA or SNr while D2R-SPNs project to the ventral pallidum (VP); VP neurons then innervate the STN, which in turn connects to the SNr (46). Such a clear-cut segregation, however, is less true in the NAc, where a significant proportion of D1R SPNs project to the VP (32, 47, 48).

D1R- and D2R-SPNs of the NAc were shown to play contrasted roles in modulating behavior, with the former proposed to drive pro-reward/approach responses and the latter considered as inhibiting these responses (46, 49, 50), although such a simplistic view needs amendments (46, 51–53). Indirect evidence argues for similar opposing roles of D1R-SPNs and D2R-SPNs in modulating social behavior. Optogenetic stimulation of NAc D1R-SPNs in typically behaving mice increases social approach (13, 54); social encounter with a female mouse activates VTA-projecting D1R-SPNs in the NAc of male mice (48). In the chronic social defeat stress (SDS) model of depression, social avoidance strongly correlated with D2R-SPN excitability and stimulating these neurons in the NAc of naïve mice produced social avoidance following subthreshold SDS. In contrast, repeated optogenetic stimulation of D1R-SPNs in the NAc restored social interaction in depressive-like mice (32). In *Drd1-Cre* mice, manipulating the expression of candidate genes for autism or schizophrenia in striatal SPNs has provided further evidence for their role in controlling social behavior. Deletion of the schizophrenia candidate gene *Erbb4* simultaneously in both populations of NAc SPNs produced social impairment (55). Knocking down Neuroligin-2 in D1R-SPNs led to a deficit in social interaction and dominance, while such manipulation in D2R-SPNs resulted in increased defensive response to social aggression (56). Suppressing endocannabinoid 2-arachidonoylglycerol (2-AG) signaling in D1R-SPNs also blunted sociability, an effect mimicked by local ablation in the DS (57). Finally, knocking down the *Shank3* gene in NAc D1R-SPNs impaired social preference (54). Together, these studies point to D1R-SPNs, notably in the NAc, as driving pro-social behavior, whereas activation of D2R-SPNs would rather promote social avoidance.

The present study aimed at exploring the respective roles of D1R-SPNs and D2R-SPNs of the NAc in driving social behavior. First, we selectively ablated each of these neuronal populations using an inducible diphtheria toxin receptor (DTR)-mediated cell targeting strategy (49, 58). Social behavior assessment in ablated mice was completed by an evaluation of repetitive/perseverative behavior, motor function and anxiety levels, most often altered in neuropsychiatric disorders together with the social repertoire. We reveal a severe social behavior deficit following ablation of D1R-SPNs in the NAc, but not dorsal striatum (DS), while ablation of D2R-SPNs in the NAc facilitated social interactions. We show that administration of pharmacological compounds repressing D2R-SPN activity relieves behavioral deficits in mice bearing D1R-SPN ablation in the NAc and concomitantly represses gene expression in the ventral pallidum (VP). Finally, we evidence that long, but not short, optogenetic stimulation of NAc D2R-SPNs induces a deficit in social interaction and increases neuronal transcription in the VP and lateral hypothalamus (LH); furthermore, pharmacological inhibition of D2R-SPNs prevented optogenetically-induced social deficit in mice.

## METHODS AND MATERIAL

### Animal care

All experimental procedures were conducted in accordance with the European Communities Council Directive 2010/63/EU and the Institutional Animal Care Committee guidelines. They were approved by the Local Ethical Committees (Pôle Santé ULB, Bruxelles). Animal were housed in group (2-4 mice per cage) and maintained on a 12h light-dark cycle (light on at 08.00) at a controlled temperature of 21°C. Food and water were available *ad libitum*, except otherwise stated.

### Subjects

Heterozygous C57BL/6 *Drd1a-Cre^+/-^ EY262* strain mice (http://www.gensat.org; Gong et al, 2007) were crossed with homozygous *C57BL/6 iDTR^+/+^* mice (59). *Drd1a-Cre^+/-^ iDTR^+/^*^-^ mice expressed the diphtheria toxin receptor in D1R-SPNs neurons (58). *Drd1a-Cre^-/-^ iDTR^+/-^* mice were used as controls. Similarly, heterozygous C57BL/6 *Drd2-Cre^+/-^ ER44* strain mice [http://www.gensat.org; (60)] were crossed with homozygous *C57BL/6 iDTR^+/+^* mice to obtain *Drd2-Cre^+/-^ iDTR^+/^*^-^ mice expressing the diphtheria toxin receptor in D2R-SPNs (61). *Drd2-Cre^-/-^ iDTR^+/-^* mice were used as controls. Experiments in Fig.1 and 2 were performed in male mice only; experiments in Fig. 3, 4 and 5 were performed in equivalent numbers of male and female mice. For the social interaction and 3-chamber tests, WT gender- and age-matched C57BL/6 mice were used as naïve interactor mice. Animals were aged 12-13 weeks at the beginning of behavioral experiments. See genotyping details in supplement.

**Figure 1.**
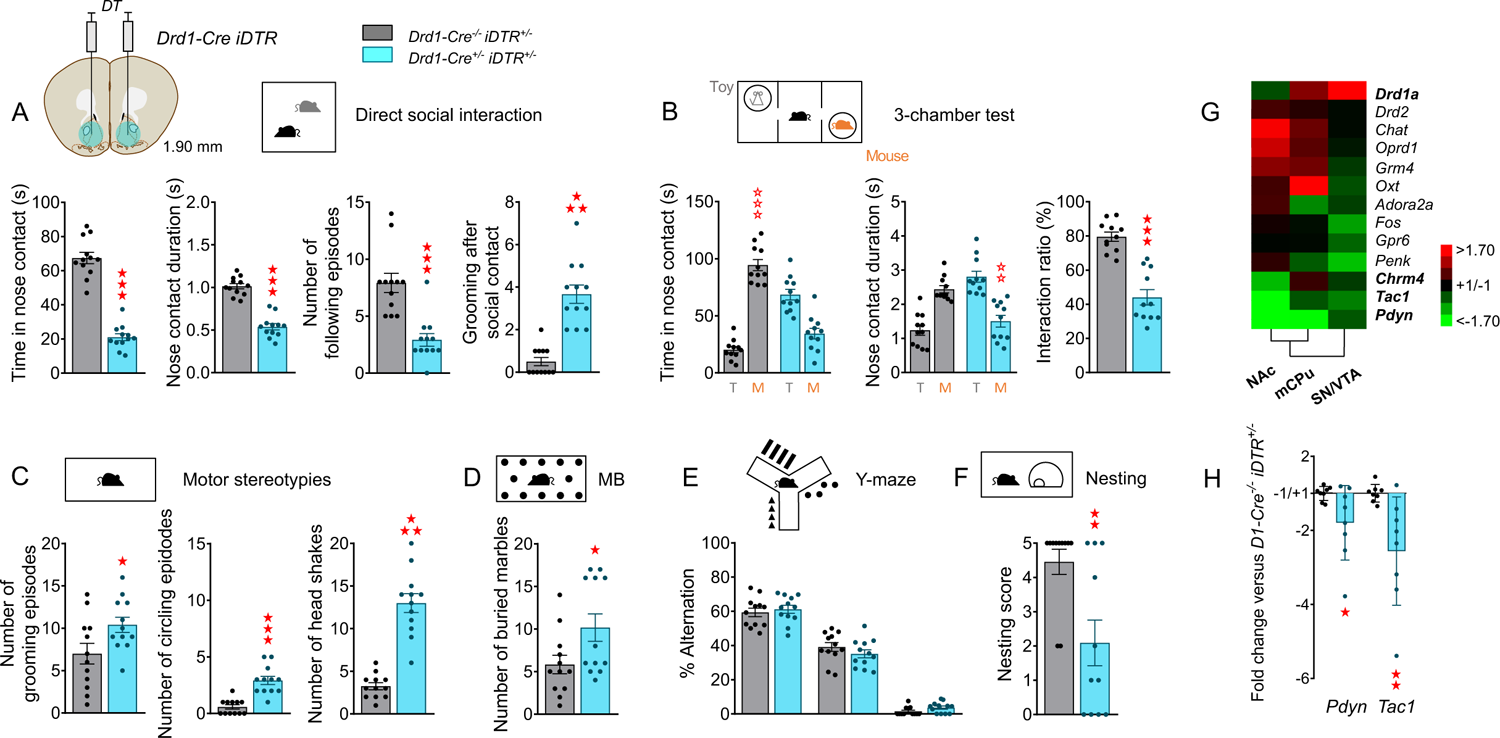
Behavioral and transcriptional consequences of ablating D1R-SPNs in the nucleus accumbens. **(A)** Experimental Design: adult *Drd1-Cre^+/-^ iDTR^+/-^* mice and *Drd1-Cre^-/-^ iDTR^+/-^* controls (n=11-14 per group) were injected stereotaxically with diphteria toxin (DT) in the NAc. Behavioral assays were performed successively between D21 and D40. **(B)** In the social interaction test, mice bearing D1R-SPN ablation (*Drd1-Cre^+/-^ iDTR^+/-^*) in the NAc displayed shorter time spent in nose contact (*Ablation: F_1,22_=141.4, p<0.0001)*, reduced duration of these contacts (*F_1,22_=96.9, p<0.0001)* and number of following episodes (*F_1,22_=24.7, p<0.0001)*, and groomed more often immediately after a social contact (*F_1,22_=44.6, p<0.0001)* compared to sham controls (*Drd1-Cre^+/-^ iDTR^+/+^*). **(C)** In the 3-chamber test, mice with NAc D1R-SPN ablation preferred to stay longer in nose contact with the toy rather than the mouse (*Stimulus x Ablation effect: F_1,20_= 40.9, p<0.0001*) and made longer-lasting contacts with the toy (*S x A: F_1,22_= 18.21, p<0.001)*, resulting in a decrease in their interaction ratio compared to sham mice (*F_1,20_=45.8, p<0.0001)*. **(D)** Compared to sham mice, lesioned mice displayed more frequent grooming (*F_1,22_=5.1, p<0.05)*, circling *(F_1,22_=32.9, p<0.0001)* and head shaking episodes (*F_1,22_=68.8, p<0.0001)*, **(E)** and buried more marbles in marble burying test (*H_1,24_=4.1, p<0.05)*, **(F)** without displaying perseverative same arm entries in the Y-maze. **(G)** Mice with NAc D1R-SPN ablation displayed lower nest building score than controls (*H_1,22_=9.5, p<0.01)*. **(H)** Gene markers of D1R- (in bold) and not D2R-SPNs were down-regulated in the NAc of ablated mice, **(I)** notably the expression of the main markers *Pdyn* and *Tac1* (see Table S1). Behavioral and transcriptional results (fold change compared to sham controls) are shown as scatter plots and mean ± sem. Solid stars: ablation effect (one-way or Kruskal-Wallis ANOVA), open stars: ablation x stimulus interaction (two-way ANOVA, mouse versus object comparison). One symbol: p<0.05, two symbols: p<0.01; three symbols: p<0.001. More behavioral parameters in Fig S1. 3-Ch: 3-chamber test, AAR: alternate arm returns, D: day, EPM: elevated plus-maze, M: mouse, MB: marble burying, mDS: medio dorsal striatum, NAc: nucleus accumbens, Nest: Nesting, NSF: novelty-suppressed feeding, RR: rotarod, SAR: same arm returns, SI: social interaction, SN/VTA: substantia nigra/ventral tegmental area, SPA: spontaneous alternation, Stereo: stereotypies, T: toy, Y-M: Y-maze.

**Figure 2.**
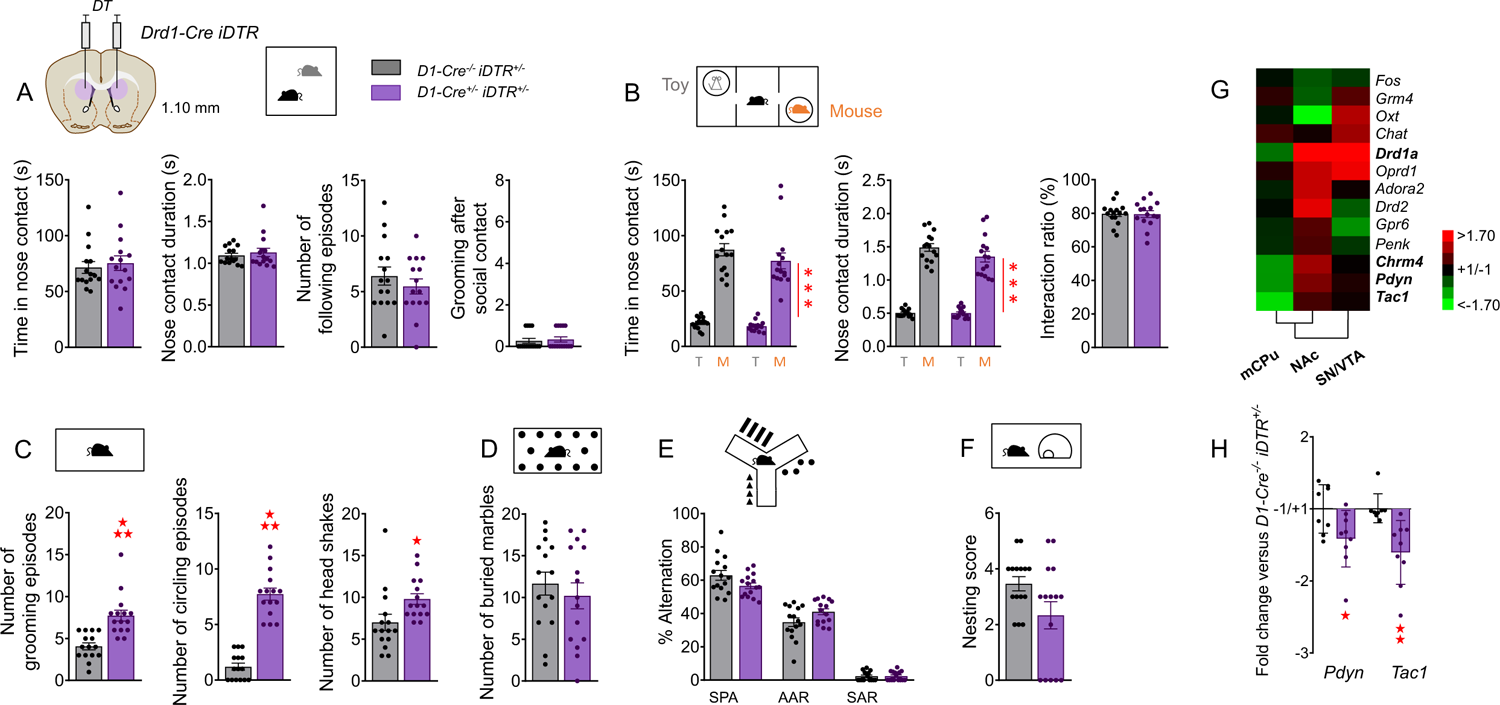
Behavioral and transcriptional consequences of ablating D1R-SPNs in the dorsal striatum. **(A)** Experimental design. Adult *Drd1-Cre^+/-^ iDTR^+/-^* mice and *D1-Cre^-/-^ iDTR^+/-^* controls (n=15 per group) were injected stereotaxically with diphteria toxin (DT) in the mDS. Behavioral assays were performed successively between D21 and D40. **(B)** Mice with D1R-SPN ablation (*Drd1-Cre^+/-^ iDTR^+/-^*) in the mDS showed similar social interaction **(C)** and social preference than sham controls (*D1-Cre^+/-^ iDTR^+/+^*). **(D)** D1R-SPN ablation in the mDS increased spontaneous stereotypic grooming (*F_1,28_=22.4, p<0.0001)* and circling episodes (*F_1,28_=107.6, p<0.0001)*, as well as head shaking (*F_1,28_=5.7, p<0.05)*, **(E)** but did not burry more marbles in the marble burying test **(F)** nor exhibited perseverative same arm entries in the Y-maze. **(G)** D1R-SPN ablation in the mDS failed to affect significantly nest building. **(H)** Gene markers of D1R-SPNs (in bold) were down-regulated in the mDS of ablated mice, **(I)** notably the expression of the main markers *Pdyn* and *Tac1* (see Table S2). Behavioral and transcriptional results (fold change compared to sham controls) are shown as scatter plots and mean ± sem. Solid stars: ablation effect (one-way or Kruskal-Wallis ANOVA), asterisks: stimulus effect (two-way ANOVA). One symbol: p<0.05, two symbols: p<0.01; three symbols: p<0.001. More behavioral parameters in Fig S2. 3-Ch: 3-chamber test, AAR: alternate arm returns, D: day, EPM: elevated plus-maze, M: mouse, MB: marble burying, mDS: medio dorsal striatum, NAc: nucleus accumbens, Nest: Nesting, NSF: novelty-suppressed feeding, RR: rotarod, SAR: same arm returns, SI: social interaction, SN/VTA: substantia nigra/ventral tegmental area, SPA: spontaneous alternation, Stereo: stereotypies, T: toy, Y-M: Y-maze.

**Figure 3.**
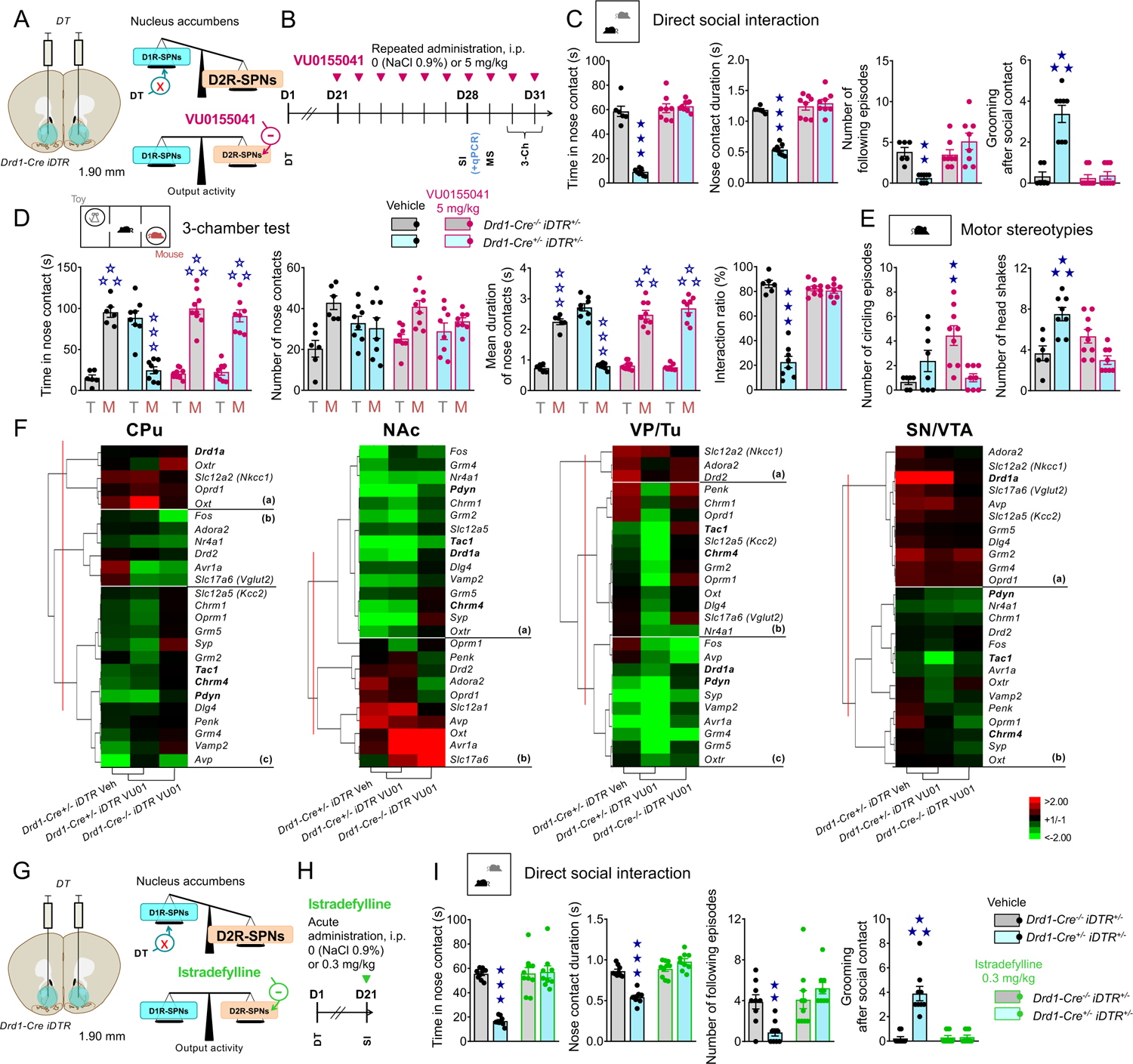
Behavioral and transcriptional consequences of D2R-SPN ablation in the nucleus accumbens. **(A)** Experimental design. Adult *D2-Cre^+/-^ iDTR^+/-^* mice and *D2-Cre^-/-^ iDTR^+/-^* controls (n=10 per group) were injected stereotaxically with diphteria toxin (DT) in the NAc. Behavioral assays were performed successively between D21 and D40. **(B)** In the social interaction test, mice bearing D2R-SPN ablation (*Drd2-Cre^+/-^ iDTR^+/-^*) in the NAc spent more time in nose contact (*F_1,17_=74.8, p<0.0001)* and made longer nose contacts (*F_1,17_=6.5, p<0.05)* with the unfamiliar conspecific; they displayed more frequent following episodes (*F_1,17_=37.0, p<0.0001)* and groomed less after a social contact (*F_1,17_=14.3, p<0.01)* than sham controls (*Drd2-Cre^+/-^ iDTR^+/+^*). **(C)** In the 3-chamber test, mice with NAc D2R-SPN ablation showed exacerbated social preference, as shown by longer time spent in nose contact (*S x A: F_1,17_=39.3, p<0.0001)* and longer mean duration (*S x A: F_1,17_=9.7, p<0.01)* of these contacts with the mouse over the toy than sham controls, resulting in increased interaction ratio (*F_1,17_=16.7, p<0.001)*. **(D)** D2R-SPN ablation in the NAc resulted in increased number of spontaneous circling episodes (*F_1,17_=20.7, p<0.001)*, **(E)** decreased number of buried marbles in the marble burying test (*H_1,19_=11.7, p<0.001)*. **(F)** Mice with NAc D2R-SPN ablation made more alternate arm returns (AAR, *F_1,17_=14.7, p<0.01*) and less same arm returns (SAR, *F_1,17_=30.6, p<0.0001*) in the Y-maze and **(G)** displayed similar nesting scores as sham controls. **(H)** Gene markers of D2R-SPNs (in bold) were down-regulated in the NAc of ablated mice, as well as in the mDS, likely along the track of the injection canula. **(I)** The two main gene markers of D2R-SPNs *Penk* and *Adora2a* were down-regulated in the NAc of D2R-SPN ablated mice, as well as the *Oprd1* marker gene of CINs (see Table S3). Behavioral and transcriptional results (fold change compared to sham controls) are shown as scatter plots and mean ± sem. Solid stars: ablation effect (one-way or Kruskal-Wallis ANOVA) open stars: ablation x stimulus interaction (two-way ANOVA, mouse versus object comparison), (a): ablation x stimulus interaction (comparison with sham mouse condition, p<0.001). One symbol: p<0.05, two symbols: p<0.01; three symbols: p<0.001. More behavioral parameters in Fig S3. 3-Ch: 3-chamber test, AAR: alternate arm returns, D: day, EPM: elevated plus-maze, M: mouse, MB: marble burying, mDS: medio dorsal striatum, NAc: nucleus accumbens, Nest: Nesting, NSF: novelty-suppressed feeding, RR: rotarod, SAR: same arm returns, SI: social interaction, SN/VTA: substantia nigra/ventral tegmental area, SPA: spontaneous alternation, Stereo: stereotypies, T: toy, Y-M: Y-maze.

**Figure 4.**
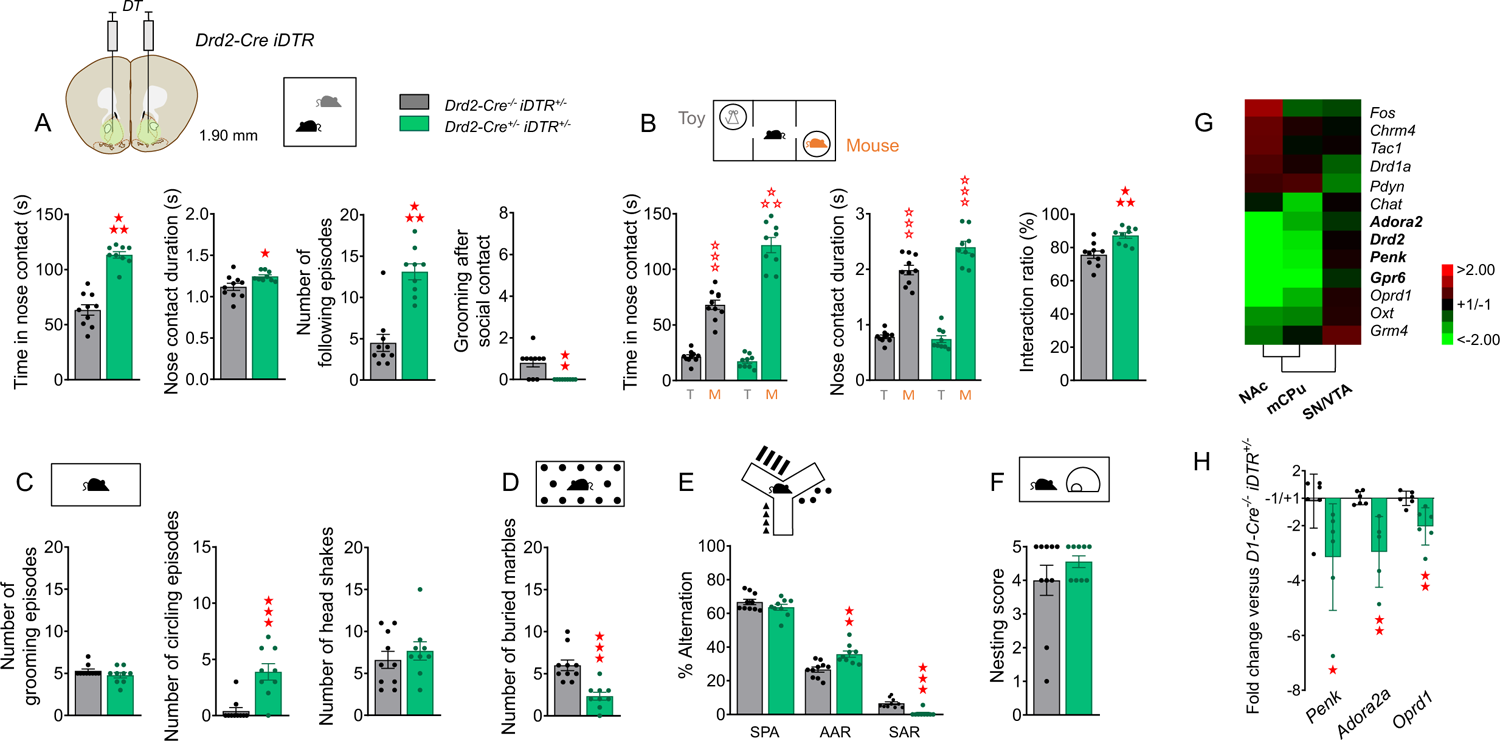
Behavioral and transcriptional consequences of D2R-SPN pharmacological inhibition in mice bearing D1R-SPN ablation in the nucleus accumbens. **(A)** Experimental design. Mice bearing D1R-SPN ablation in the NAc and controls were treated chronically with the mGlu4 PAM VU0155041, to rebalance D1R/D2R-SPN output activity balance. They received VU0155041 (5 mg/kg, i.p.) or vehicle (NaCl, 0.9%) daily from D21 after surgery to D31. Behavioral testing was performed from D28 to D31 (cohort 1, n=6-8 per group). In cohort 2 (n=6 per group), mice were euthanized 45 min after direct social interaction (D28) and brain samples collected for qRT-PCR analysis. **(B)** Chronic VU0155041 administration normalized social interaction parameters in NAc D1R-SPN ablated mice, as illustrated by rescued time spent in (*Treatment x Ablation: F_1,26_=84.8, p<0.0001*) and mean duration of (*T x A: F_1,26_=43.7, p<0.0001*) nose contacts, number of following episodes (*T x A: F_1,26_=12.8, p<0.0001*) and grooming after a social contact (*T x A: F_1,26_=28.0, p<0.0001*). **(C)** Similarly, in the 3-chamber test, chronic facilitation of mGlu4 activity restored preference for spending more time (*S x T x A: F_1,27_=56.3, p<0.0001*) and making longer nose contacts (*S x T x A: F_1,27_=146.4, p<0.05*) with the mouse over the toy in ablated mice, resulting in normalized interaction ratio (*T x A: F_1,26_=4.3, p<0.0001*). **(D)** Finally, VU0155041 treatment suppressed spontaneous circling episodes (*T x A: F_1,27_=14.2, p<0.001*) and head shakes (*T x A: F_1,27_=23.9, p<0.0001*) in mice bearing D1R-SPN ablation. **(E)** Gene expression was not restored upon chronic VU0155041 administration in the NAc of NAc D1R-SPN ablated mice, but was widely repressed in the VP of these animals compared to saline-treated sham controls (see Table S4). **(F)** Experimental design. Mice bearing D1R-SPN ablation in the NAc and controls were treated acutely with the A2a adenosine receptor antagonist istradefylline, to rebalance D1R/D2R-SPN output activity balance. Istradefylline (0.3 mg/kg, i.p.) or vehicle (NaCl 0.9%) were administered acutely (n=9-10 per group) 30 min before testing (D21 after surgery). **(G)** Acute istradefylline normalized the time spent in (*T x A: F_1,33_=30.2, p<0.0001*) and duration of (*T x A: F_1,33_=48.9, p<0.0001*) nose contacts, as well the number of following episodes (*T x A: F_1,33_=9.5, p<0.01*) and grooming episodes after social contact (*T x A: F_1,33_=32.9, p<0.0001*) in NAc D1R-SPN ablated mice. Behavioral results are shown as scatter plots and mean ± sem. Solid stars: ablation x treatment interaction (two-way ANOVA), open stars: ablation x stimulus interaction (three-way ANOVA, mouse versus object comparison). One symbol: p<0.05, two symbols: p<0.01; three symbols: p<0.001. More behavioral parameters in Fig S4. 3-Ch: 3-chamber test, D: day, DT: diphtheria toxin, M: mouse, mDS: medio-dorsal striatum, MS: motor stereotypies, NAc: nucleus accumbens, SI: social interaction, SN/VTA: substantia nigra/ventral tegmental area, T: toy, VP/Tu: ventral pallidum/olfactory tubercle.

**Figure 5.**
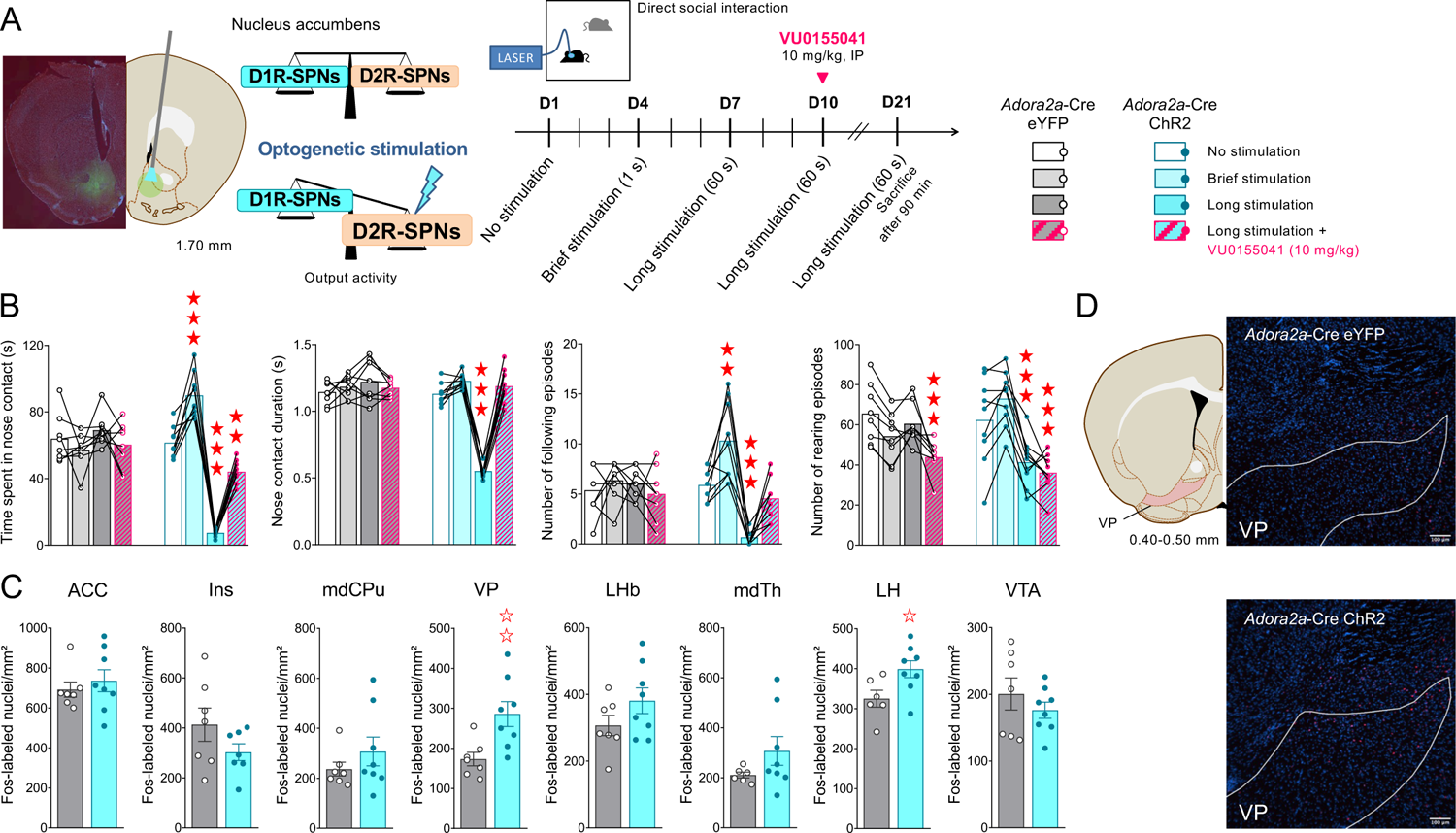
Effects of optical stimulation of D2R-SPN in the nucleus accumbens on social interaction and Fos expression in *Adora2a-cre* mice. **(A)** Experimental design. *Adora2a-cre* mice (n=8-9) were injected in the NAc with AAV5 EF1a-DIO-hChR2(H134R)-eYFP containing a cre-dependent channelrhodopsin (ChR2) or eYFP and direct optical stimulation was applied to mimic excessive D2R-SPN activity. Social interaction of *Adora2a-cre-*ChR2 mice and *Adora2a-cre-*YFP controls was first assess in the absence of light (D1), then under short stimulation conditions (light on 1 s, D4) and under long stimulation conditions (light on 60 s, D7). On D10, all mice received VU0155041 (10 mg/kg) 30 min before being tested for social interaction under prolonged stimulation conditions. After a week drug-free, all mice were again submitted to a social interaction test under prolonged stimulation conditions (D21) and sacrificed 90 min after for Fos immunostaining experiments. **(B)** Brief optical D2R-SPN stimulation in the NAc facilitated direct social interaction, as shown by increased time spent in nose contact and number of following episodes in *Adora2a-cre-*ChR2 mice. In contrast, prolonged stimulation markedly decreased social interaction, as evidenced by reduced time spent in and mean duration of nose contacts, as well as reduced number of following episodes in *Adora2a-cre-*ChR2 mice. Their number of rearing episodes was also reduced under these conditions. Finally, chronic VU0155041 treatment opposed the deleterious effects of prolonged stimulation on social interaction parameters, increasing the time spent in nose contacts (*Viral construct x Condition: F_3,45_*_=_*42.8, p<0.0001*) and restoring nose contact duration (*V x C: F_3,45_*_=_*80.4, p<0.0001)* and number of following episodes (*V x C: F_3,45_*_=_*11.0, p<0.0001*), without normalizing the number of rearing episodes. **(C)** We detected an increase in Fos expression in two output regions of the NAc, the VP and LH. **(D)** Representative microphotographs showing Fos positive nuclei in the VP of *Adora2a-cre-*YFP (upper panel) and *Adora2a-cre-*ChR2 (lower panel) after prolonged NAc D2R-SPN optical stimulation. Magnification: x6.25, Scale bar: 100 µm. Behavioral and immunohistochemical results are shown as scatter plots and mean ± sem. Solid stars: optical stimulation effect (two-way ANOVA, compared to no light condition in *Adora2a-cre-*YFP mice), open stars: optical stimulation effect stimulus interaction (Student t test, compared to prolonged stimulation condition in *Adora2a-cre-*YFP mice). One symbol: p<0.05, two symbols: p<0.01; three symbols: p<0.001. More behavioral parameters in Fig S5. ACC: anterior cingulate cortex, Ins: insular cortex, LH: lateral hypothalamus, LHb: lateral habenula, mDS: medio-dorsal striatum, mdTh: medio-dorsal thalamus, VP: ventral pallidum, VTA: ventral tegmental area.

### Toxin ablation and optogenetic manipulation

#### Ablation

*Drd1a-Cre^+/-^ iDTR^+/-^* and *Drd1a-Cre^-/-^ iDTR^+/-^* or *Drd2-Cre^+/-^ iDTR^+/-^* and *Drd2-Cre^-/-^ iDTR^+/-^* mice were deeply anesthetized at the age of 8 weeks and placed on a stereotaxic frame, as described previously (49,58,61). The stereotaxic injection of diphtheria toxin (DT; Sigma-Aldrich; 100 pg/µl; diluted in 0.01M PBS) was performed with a blunt needle in the striatum. For ablation in the NAc, DT was administered bilaterally in two symmetrical injections of 0.2 µl at a slow rate of 0.1 µl/min and the needle was removed 20 min after the end of the injection to ensure optimal diffusion. The following coordinates (adapted from atlas of Paxinos and Franklin, 2001, bregma as reference) were used: NAc - anterior +1.9 mm; lateral ±1.2 mm; ventral −4.9 mm. For ablation in the DS, DT was administered bilaterally in 2 symmetrical injections of 1µl at a slow rate of 0.25 µl/min and the needle was removed 15 min after the end of the injection to ensure optimal diffusion. The following coordinates were used: DS: anterior + 0.9mm; lateral ±1.5 mm; ventral ± 3.0 mm and anterior + 0.5mm; lateral ±1.8 mm; ventral ±3.0 mm. Control of D1R-SPN ablation in the NAc was performed by assessing specific SPN markers using qRT-PCR. After stereotaxic injections, mice recovered from the surgery for two weeks.

#### Optogenetics

We injected 0,5 µL of either AAV5 EF1a-DIO-hChR2(H134R)-eYFP (from UNC) or AAV5 EF1a-DIO-eYFP (from UNC) at the coordinates: AP+1.85, ML+-0.95, DV-4.55. Optics fibers (0.22 NA, Ø200 µm Core fibers (ThorLabs FB200UEA) and 1.25mm Multimode LC|PC SS Ø231 µm Ferrules (ThorLabs SFLC230) were implanted during the same surgery with an angle of 10° using the coordinates: AP+1.85, ML+-1.60, DV-3.55. Brief optical stimulation was performed as follows: 473 nm; frequency of 40 Hz; 12 ms pulses; 5 mW at the output of the patch-cord; 1 min OFF followed by 9 cycles (1 sec ON / 59 sec OFF) over 9 min. Prolonged optical stimulation was performed as follows: 473 nm; frequency of 40 Hz; 12 ms pulses; 5 mW at the output of the patch-cord; five cycles (1 min OFF / 1 min ON) over 10 min.

### Behavioral experiments

As described previously (62–64), social behavior was explored using the direct social interaction and three-chamber tests, stereotyped/perseverative behavior was assessed by scoring spontaneous motor stereotypies and marble burying, and by analyzing alternation patterns in the Y-maze. Motor function and coordination were measured using the nest building and accelerating rotarod tests and, finally, anxiety-like behavior was evaluated in the novelty-suppressed feeding (NFS) and elevated plus-maze tests (EPM). Detailed protocols for behavioral tests are described in Supplement. Experiments were conducted and analyzed blind to genotype and experimental condition.

### Statistics

Statistical analyses were performed using Statistica 9.0 software (StatSoft, Maisons-Alfort, France). For all comparisons, values of p<0.05 were considered as significant. Statistical significance in behavioral experiments was assessed using one to three-way analysis of variance (ablation, stimulus – mouse versus toy -, trial, session and treatment effects) followed by Newman-Keules post-hoc test or using a Kruskal Wallis ANOVA (marble burying, nesting). Significance of quantitative real-time PCR (qRT-PCR) results was assessed after transformation using a two-tailed t-test, as previously described (62, 63); an adjusted p value was calculated using Benjamini-Hochberg correction for multiple testing. Outliers over twice the standard deviation were excluded from calculations. Unsupervised clustering analysis was performed on transformed qRT-PCR data using complete linkage with correlation distance (Pearson correlation) for drug, treatment and brain region (Cluster 3.0 and Treeview software). Details for drugs and methods of quantitative real time PCR analysis, immunohistochemistry and cell counting are available in Supplement.

## RESULTS

### Ablation of D1R-SPNs in the NAc undermines social behavior, induces motor stereotypies and increases conflict anxiety

We first evaluated the consequences of D1R-SPN ablation in the NAc on social and stereotypic behavior, motor abilities and anxiety (Fig. 1A). We assessed social abilities in mice bearing D1R-expressing neuron ablation in the NAc using two tests. In the direct social interaction test (Fig. 1B), lesioned animals made less and shorter nose contacts and followed less the interactor mouse than sham mice; they groomed more often immediately after a social contact, a sign of social discomfort. In the 3-chamber social preference test (Fig. 1C), lesioned animals showed decreased preference to interact with the living interactor compared to the toy.

As regards stereotypic and repetitive behavior, D1R SPN-lesioned mice displayed more frequent spontaneous motor stereotypies (Fig. 1D) and buried more marbles (Fig. 1E) than sham controls. In the Y-maze (Fig. 1F), however, these mice did not display increased perseverative same arm entries. We then assessed motor function in D1R-SPN-ablated mice. Nest building (Fig. 1G), a social-related motor response (65), was impaired in ablated mice. Finally, NAc D1R-SPN ablation facilitated motor skill learning (Fig S2E), reduced the latency to feed in the NSF test (Fig S2F) that assesses approach/avoidance behavior (66) and explored less the open arms in the EPM (Fig S2G) as compared to control mice.

To assess the efficiency and specificity of D1R-SPN ablation in the NAc, we used qRT-PCR and measured in the NAc, mDS and SNc/VTA the expression of 11 marker genes of striatal neural populations: D1R-expressing SPNs (*Drd1a*, *Tac1*, *Pdyn*, *Chrm4*), D2R-expressing SPNs (*Drd2*, *Adora2a*, *Penk*, *Gpr6*, *Grm4*) and cholinergic interneurons (CINs: *Chat*, *Oprd1*), as well as the expression of *Fos*, as a marker of neuronal activity, and *Oxt* (coding for oxytocin) as a marker of social behavior, 45 min after a direct social interaction (Fig. 1H and Table S2). Ablation of NAc D1R-SPNs was associated with a significant decrease in the expression of *Tac1* and *Pdyn*, the two main marker genes of D1R-expressing SPNs, in this structure, confirming D1R-SPN ablation (Fig. 1I). In contrast, the expression of two marker genes of D2R-SPNs, *Drd2* and *Grm4*, and markers of CINs *Oprd1* and *Chat* were increased in the NAc. In the mDS, D1R-SPN ablation was associated with decreased expression of *Pdyn* but increased expression of *Drd1a*. Finally, in the SN/VTA, *Drd1a* expression was also found up-regulated, while *Penk* expression was down-regulated. These additional regulations may reflect adaptative modulation of gene expression as a compensation for NAc D1R-SPN ablation, while decreased *Pdyn* in the mDS may reflect some partial D1R-SPN deletion along the injection track.

Therefore, selective ablation of D1R-SPNs in the NAc, evidenced by reduced expression of specific gene markers *Tac1* and *Pdyn* in this region, resulted in social interaction deficit, stereotypic behavior, facilitated motor skill learning and increased anxiety levels in EPM.

### Ablation of D1R-SPNs in the DS induces stereotypies, impairs motor learning and decreases anxiety without impacting social behavior

We then assessed the effects of D1R-SPN ablation in the DS on social and stereotypic behavior, motor abilities and anxiety to evaluate the specificity of behavioral impairments detected following NAc D1R-SPN ablation (Fig. 2A). In the direct social interaction test (Fig. 2B), selective lesion of D1R-SPNs had no detectable impact on the measured parameters. Consistent with this, mice bearing a deletion of D1R-SPNs in the DS showed similar preference to interact with a mouse in the 3-chamber test as sham controls (Fig. 2C). In contrast, when observed for 10 min under mildly anxiogenic conditions, these animals displayed more frequent motor stereotypies than control mice (Fig. 2D). However, no significant effect of D1R-SPNs ablation was observed in the marble burying test (Fig. 2E), on the pattern of exploration in the Y-maze test (Fig. 2F) nor on nest building performance (Fig. 2G). Nonetheless, this lesion produced a severe impairment in learning a motor skill on the accelerating rotarod (Fig. S3E), as previously shown (58). Finally, mice bearing D1R-SPN ablation in the DS displayed shorter latencies to eat in the novelty-suppressed feeding test (Fig. S3F) and, in the EPM test (Fig. S3G), explored more the open arms compared to sham controls, suggesting reduced levels of anxiety.

At transcriptional level (Fig. 2J-K, Table S3), deletion of DS D1R-SPNs was consistently associated with a significant decrease in *Pdyn* and *Tac1* expression in the mDS. In the NAc, the expression of *Drd1a*, *Oprd1*, *Adora2a* and *Pdyn* were found upregulated; in the SNc/VTA, *Drd1a* and *Oprd1* expression were increased. These results indicate that D1R-SPN ablation in the DS did not spread to the NAc (where no decrease in D1R-SPN marker genes was detected), and that adaptations in gene expression developed in regions connected to lesioned region.

Together, these data indicate that the selective ablation of D1R-SPNs in the DS, confirmed by reduced expression of *Pdyn* and *Tac1* in this region, did not alter social behavior in mice, but increased spontaneous stereotyped behaviors, impaired motor skill learning and decreased anxiety-like behavior.

### Ablation of D2R-SPNs in the NAc increases social approach and impairs motor learning

To address the question of the respective contributions of D2R-SPNs versus D1R-SPNs of the NAc in controlling behavior, we subjected mice bearing D2R-SPN ablation in the NAc to the same behavioral assays as previously (Fig. 3A). Remarkably, D2R-SPN lesion increased social interaction while reducing grooming after social contact in the direct social interaction test (Fig. 3B). Accordingly, animals with D2R SPN ablation displayed greater preference for interacting with a mouse over a toy in the 3-chamber test (Fig. 3C). In ablated mice, the number of spontaneous circling episodes was significantly increased (Fig. 3D); in contrast, these mice buried less marbles (Fig. 3E). When exploring the Y-maze, D2R-SPN-lesioned mice made more frequent alternations between two arms, but less perseverative same arm entries than sham controls (Fig. 3F). These mice showed normal performance in building a nest (Fig. 3G) but exhibited impaired motor learning on the accelerating rotarod (Fig. S4E). Finally, ablation of D2R-SPNs in the NAc had no impact on anxiety levels in the novelty-suppressed feeding (Fig. S4F) and elevated plus-maze (Fig. S4G) tests.

At transcriptional level (Fig. 3H-I, Table S4), D2R-SPN ablation in the NAc was associated with a marked decrease in the expression of the D2R-SPN marker genes *Penk*, *Adora2a, Drd2* and *Gpr6*, as well as in *Oxt* and the CIN marker gene *Oprd1*; *Fos* expression was increased. In the mDS, the expression of *Penk*, *Adora2a*, *Gpr6* and *Drd2* were found down-regulated, likely along the track of the injection canula, as well as the expression of CIN markers *Chat* and *Oprd1*. In the VTA, the expression of *Grm4* was upregulated, showing little impact of D2R-SPN ablation.

Thus, D2R-SPN lesion in the NAc, likely associated with CIN lesion, as evidenced by reduced *Penk*, *Adora2a* and *Oprd1* expression in the NAc, facilitated social interaction and preference, increased circling behavior but decreased marble burying and perseveration in the Y-maze, and hindered motor skill learning, with no effect on anxiety-like behavior.

### Pharmacological inhibition of D2R-SPN activity relieves behavioral deficits and represses VP gene expression in mice with D1R-SPN ablation in the NAc

As we hypothesized that a balance between D1R-SPN and D2R-SPN activity in the NAc controls social and stereotypic behaviors, we tested whether administering pharmacological compounds that dampen D2R-SPN activity may alleviate the behavioral impairments of mice bearing a D1R-SPN ablation in this region.

We chronically treated mice bearing an ablation of D1R-SPN in the NAc with the positive allosteric modulator (PAM) of the mGlu4 receptor VU0155041 (5 mg/kg, i.p.) (Fig. 4A), a repressor of D2R-SPN activity that relieved autistic-like symptoms in the *Oprm1* mouse model of ASD (67). We then assessed autism-sensitive behaviors and gene expression. Chronic VU0155041 administration normalized direct social interaction in NAc D1R-SPN ablated mice (Fig. 4B). In the three-chamber test (Fig. 4C), mGlu4 PAM administration restored the preference of lesioned mice for exploring with the living interactor. Finally, facilitating mGlu4 function normalized the number of circling episodes and head shakes in D1R-SPN lesioned animals (Fig. 4D, see more parameters in Fig S5). Chronic mGlu4 PAM treatment was thus able to relieve the deleterious behavioral effects of D1R-SPN ablation in the NAc.

To identify molecular substrates underlying the beneficial effects of chronic VU0155041 treatment in mice bearing a selective ablation of D1R-SPNs in the NAc, we assessed the expression of 24 genes, including marker genes of D1R-SPNs (*Drd1a*, *Tac1*, *Pdyn*, *Chrm4*), D2R-SPNs (*Drd2*, *Adora2*, *Penk*, *Gpr6*, *Grm4*), D2R-SPNs (*Chrm1, Grm2, Grm5, Nr4a1, Oprm1*) or CINs (*Chat*, *Oprd1*), genes of the oxytocin/vasopressin system (*Oxt, Oxtr, Avp, Avr1a*), known to play a key role in controlling social behavior (68, 69), genes coding for chloride and glutamate transporters (*Slc12a2, Slc12a5, Slc17a6), Fos* as a marker of neuronal transcription and finally *Dlg4*, *Syp* and *Vamp2* as marker genes of synaptic function, 45 min after a direct social interaction in the NAc, mDS, ventral pallidum/olfactory tubercle (VP/Tu) and SNc/VTA in mGlu4 PAM versus vehicle treated mice. As shown by cluster analysis (Fig. 4E and Table S5), chronic VU0155041 had little effect on gene expression in the DS and VTA/SNc. This treatment did not either modify the pattern of gene expression associated with D1R-SPN ablation in the NAc of *Drd1^-^Cre^+/-^* mice (cluster a). In contrast, in the VP/Tu, VU0155041 triggered a marked down-regulation of gene expression (clusters b,c), in addition to that produced by NAc D1R-SPN ablation (cluster c). Together, these results point to VP/Tu region as a key target for transcriptional effects of VU0155041 in D1R-SPN ablated mice.

To confirm our hypothesis, we evaluated the behavioral effects of another pharmacological repressor of D2R-SPN activity, the adenosine A2a receptor antagonist istradefylline (70), in NAc D1R-SPN-ablated mice (Fig. 4F). Direct social interaction was assessed after an acute istradefylline injection (0.3 mg/kg). At this dose, istradefylline completely restored direct social interaction parameters (Fig. 4G, more parameters in Fig S5). Therefore, istradefylline was able to alleviate social interaction deficits in mice bearing NAc D1R-SPN ablation.

Thus, pharmacological treatments that repress the activity of D2R-SPNs can relieve the deleterious effects of NAc D1R-SPN lesion on autism-sensitive behavior. Beneficial behavioral effects of the mGlu4 PAM VU0155041 in NAc D1R-SPN-ablated mice were associated with a downregulation of the expression of numerous neuronal gene markers in the VP/Tu.

### Prolonged, and not short, optical stimulation of D2R-SPN in the NAc hampers social interaction in mice

To further challenge our hypothesis and avoid nonspecific effect of pharmacological treatment, we tested whether optical stimulation of D2R-SPNs, to mimick excessive D2R-SPN output activity, may affect social behavior in mice. *Adora2a-cre* mice were injected in the NAc with AAV5-EF1-DIO-h-ChR2-YFP containing a cre-dependent channelrhodopsin (ChR2) or eYFP (Fig. 5A). Based on previous study showing differential outcome on reward/aversion (71), we compared the effects of short (1 s) versus prolonged (60 s) optical D2R-SPN stimulation on direct social interaction in NAc AAV-injected mice. We then assessed the effects of acute VU0155041 administration (10 mg/Kg, 30 min before testing) on further long D2R-SPN optical stimulation. We finally controlled optogenetic activation by Fos staining in the NAc.

In the direct social interaction test (Fig. 5B), brief optical D2R-SPN stimulation facilitated social contacts, while prolonged stimulation inhibited them in *Adora2a-cre-* ChR2 mice; prior VU0155041 treatment prevented this long stimulation-provoked deficit in social interaction (more parameters in Fig S6). Beneficial effects of VU0155041 treatment were independent from effects on vertical activity. We evaluated the neuronal consequences of long D2R-SPN stimulation by assessing Fos expression in several NAc afferences/efferences after a last social interaction test (Fig. 5C). Under these conditions, we detected an increase in Fos expression in two output regions of the NAc, the VP (Fig. 5D) and LH.

Together, our data indicate that prolonged optical stimulation of NAc D2R-SPNs impedes social interaction, an effect that can be blocked by prior administration of a mGlu4 PAM and is associated with increased neuronal activity in the VP and LH.

## DISCUSSION

Based on classical dichotomy between D1R-SPNs and D2R-SPNs in driving reward versus aversion (46, 49, 50, 53), activation of D1R-SPN has been proposed to trigger social approach whilst activation of D2R-SPN would lead to social withdrawal. In the present study, we evidenced a severe deficit in social interaction and preference following selective ablation of D1R-SPNs in the NAc. When D1R-SPN ablation targeted the DS, no social deficit was detected. These findings are consistent with a prosocial role of D1R-SPNs in the NAc, in agreement with their activation (VTA-projecting D1R-SPNs) during social encounter (48) and the effects of their optogenetic stimulation in typically behaving (13, 54) or depressive-like (32) mice. In contrast, *Shank3* knockdown in NAc D1R-SPN increased their excitability and concomitantly reduced social preference in mice (54), suggesting that excessive D1R-SPN activity may also lead to social avoidance, possibly by involving VP-projecting populations (48). However, increased D1R-SPN excitability in *Shank3* mutant mice can co-exist with decreased output activity, as seen in mouse models of depression (32). Deleting diacylglycerol lipase *α* gene (*Dglα*) from D1R-SPNs increased their spontaneous activity; when this deletion was restricted to the mDS, but not the NAc, social preference was impaired (57), as observed following D1R stimulation (72), indicating that a role of DS D1R-SPNs in controlling social behavior should not be discarded.

Mirroring the effects of D1R-SPN ablation, D2R-SPN ablation in the NAc facilitated social behavior. Consistent with our data, mice lacking D2R, which inhibits the activity of D2R-SPNs, showed deficient social behavior (72). Spontaneous activity of D2R-SPN was increased in the NAc of depressive-like mice correlating negatively with social interaction (32). These elements argue for a tonic repressing role of D2R-SPNs, notably of the NAc, on social behavior. Not only NAc D2R-SPNs were ablated in our study, however, as revealed by concomitant decrease in D2R-SPN and CIN gene marker expression. Indeed, to drive iDTR expression, we used a *Drd2* promoter, expressed in both striatal D2R-SPNs and CINs (73, 74). CINs have also been implicated in the control of social behavior. Total striatal ablation of CINs exacerbated social behavior in mice (75). In contrast, DS-restricted CIN deletion reduced social interaction in male mice (76). Thus, increased social behavior in NAc DT-injected *Drd2-Cre^+/-^ iDTR^+/-^* mice may have resulted, at least in part, from CIN ablation in this region. The net effect of CIN ablation on striatal SPN output activity is not fully elucidated yet; in the DS, CINs activation by thalamic inputs biases striatal output towards enhanced responsiveness of D2R-SPNs to corticostriatal inputs (77). If this applied to NAc circuitry, NAc CIN ablation would reduce D2R-SPN activity, resembling a D2R-SPN ablation; exacerbated social behavior following D2R-SPN and/or CIN ablation in the NAc may then result from reduced D2R-SPN output.

Besides social behavior, SPN ablation increased motor stereotypies. The role of SPNs in stereotypic behaviors remains elusive, as insufficient D1R-SPN output (78), excessive D2R-SPN activity or, conversely, insufficient D2R-SPN activity (79) were all reported to cause stereotypies. In contrast, complete ablation of striatal CINs did not produce such outcome (75), suggesting a limited contribution of this neuronal population. The impact of SPN ablation on social behavior was well correlated to nest building, considered a social-related motor response, but not to motor skill learning. Therefore, social deficits in NAc D1R-SPN lesioned mice were likely not a consequence of motor dysfunction, even though the crucial role of D1R-SPNs in controlling motor skills has been documented (49, 58, 80). Finally, modulation of emotional response following SPN ablation was not predictive of modifications in social behavior, making unlikely that modified levels of anxiety accounted for social behavior deficits or improvements. Globally, our data point to a dominant role of NAc D1R-SPNs in modulating anxiety-like behavior, opposing classical representation (81–83). Whether the effects of NAc D1R-SPN ablation indeed reflected deficient D1R-SPN output, however, is an intriguing question.

Previous studies have conceptualized a dualistic view of striatal outputs in which D1R-SPNs would drive reward/social approach and D2R-SPNs would trigger aversion/social avoidance. Consistent with this view, the main driver of social deficit (or excessive anxiety) in NAc D1R-SPN-ablated mice would less be the missing D1R-SPN outputs than the resulting excessive weight of D2R-SPN outputs. In support to this hypothesis, our pharmacological experiments showed that repressing D2R-SPN activity using mGlu4 PAM or A2a antagonist administration restored social behavior in mice bearing NAc D1R-SPN ablation. These results are consistent with beneficial effects of VU0155041 in the *Oprm1* mouse model of ASD (63, 67), in *Elfn2* null mice, displaying ASD-like behavioral features (84) and in morphine-abstinent mice (62), in which NAc D2R-SPN activity is excessive (85). Similarly, A2a antagonism was shown to restore social behavior in mice exposed to social stress (86). Importantly, A2a receptors being specifically expressed in the D2R-SPN of the striatum (87), these data point to an excess in D2R-SPN activity as the neuronal substrate of social deficit in NAc D1R-SPN ablated mice. Beneficial effects of mGlu4 PAM treatment in these mice were associated with a repression of gene transcription in the VP/Tu region. Such effect might appear surprising as the result of inhibiting a population of GABAergic neurons, the D2R-SPNs. However, previous studies have evidenced that sustained stimulation of D2R-SPNs induce presynaptic delta-opioid receptor-mediated LTD at D2R-SPN synapses, and increased neuronal activity in the VP (71, 88). Together, pharmacological and transcriptional data suggest a beneficial inhibitory action of mGlu4 PAM treatment at VP neurons in D1R-SPN ablated mice.

The second support to our hypothesis comes from optogenetic experiments. Upon short activation, social investigation was facilitated in *Adora2a-*Cre ChR2 mice, consistent with rewarding properties (46, 71, 89). Conversely, prolonged D2R-SPN stimulation in the NAc produced a severe deficit in social interaction, in accordance with aversive effects (71). Of note, the use of the *Adora2a* promoter allowed us to specifically target D2R-SPNs, and not CINs. Long-lasting stimulation increased neuronal activity in the VP, consistent with prior electrophysiological evidence (71, 88), and in the LH, two output regions of the NAc. As mentioned above, paradoxical increased activity in brain targets of D2R-SPN may involve enkephalin corelease, inducing presynaptic LTD (71, 88). Increased VP activity may dampen the activity of VTA DA neurons (71) whilst stimulation of VGLUT2 expressing LH neurons encodes aversive stimuli by stimulating VTA DA projections to the ventro-medial part of the NAc shell (90). Thus, the activation of both VP and LH may have contributed to support social aversion. Finally, mGlu4 facilitation, that reduces presynaptic GABA release from SPNs, prevented the deleterious effects of prolonged NAc D2R-SPN stimulation on social behavior. Together, these results indicate that excessive activity of D2R-SPNs in the NAc produces a deficit in social behavior.

Converging preclinical data suggest that NAc D1R/D2R-SPN output balance could be tilted towards excessive D2R-SPN activity in depression, autism and schizophrenia. This has been clearly evidenced in mouse models of depression, where D1R-SPNs show increased excitability and reduced output activity (32, 91), morphological atrophy (92), and decreased excitatory inputs (32, 91, 93). Accordingly, elevated D1R-SPN activity promotes resilience to depressive-like outcomes (32, 94). In contrast, D2R-SPNs show increased excitatory inputs and spine density in these models (32, 95), and D2R-SPN stimulation facilitates the development of depression-like behavior (32). In mouse models of ASD, even though experimental data consistently point to SPN dysfunction (72, 96–100), D1R/D2R-SPN balance has rarely been assessed. Synaptic inhibition on D1R- but not D2R-SPNs is decreased in *Neuroligin-3* mutant mice (80), but D1R/D2R-SPN output activity has not been directly assessed. The excitability, dendritic complexity and capacity to induce endocannabinoid-mediated long-term depression (eCB-LTD) are reduced in D2R-SPNs from *Shank3* mutant mice (79). Remarkably, blunted presynaptic LTD in D2R-SPNs could facilitate their tonic activity under sustained stimulation, as seen at D2R-SPN/VP synapses (88). Decreased eCB-LTD was also observed in NAc SPNs of *Fmr1* knockout mice (98) and DS SPNs of *neuroligin-3* mutant mice (97), although the affected SPN population remains to be identified. Finally, striatal D2R availability is increased under dopamine depletion in schizophrenia (101, 102). Blocking D2R activity with antipsychotics relieves positive symptoms (hallucinations, delusions) but fails to improve negative symptoms (anhedonia, social deficits), whose severity is instead negatively correlated with NAc D2R availability in patients (101). As D2R signaling inhibits D2R-SPN activity, facilitating this signaling selectively in the NAc could alleviate negative symptoms in schizophrenia. Preclinical assessment of D1R/D2R SPN activity balance, not performed yet, may allow challenging this hypothesis. If excessive D2R-MSN activity represented a transnosographical feature between ASD, depression and schizophrenia, this may account for shared deficit in social behavior and open novel therapeutic avenues.

There are limitations to our study. The primary focus of ablation experiments being the D1R-SPNs of the NAc, we used D1R-SPN ablation in the DS and D2R-SPN ablation in the NAc as controls, but we did not explore the role of D2R-SPNs in the DS. Similarly, in optogenetic studies, we did not assess the consequences of prolonged D1R-SPN stimulation of social behavior, which may result in social aversion (48, 54) as well as D2R-SPN stimulation. Further studies will be required to further investigate the consequences of D2R-SPN ablation in the DS and D1R-SPN stimulation in the NAc.

In conclusion, pharmacological compounds developed to repress D2R-SPN activity, such as mGlu4 PAMs or A2a antagonists, likely represent promising candidates to relieve the social deficit common to ASD, schizophrenia and depression.

## Acknowledgements

We thank Audrey Léauté, Yannick Corde, Delphine Houtteman, Laetitia Cuvelier and Souad Laghmari for technical assistance. BD and ADG are “Aspirant” FNRS and AKE is a research director of the FRS-FNRS and a WELBIO investigator.

## Funding

We acknowledge the following funding sources: JLM and JAJB - Région Centre (ARD2020 Biomédicament – GPCRAb); AKE - FNRS (grants 23587797, 33659288, 33659296), WELBIO (grant 30256053), Fondation Simone et Pierre Clerdent 2018 Prize and Fondation ULB. ADG was supported by a fellowship of the FRS-FNRS (Belgium). JLM and JAJB received support from the Institut National de la Santé et de la Recherche Médicale (Inserm), Centre National de la Recherche Scientifique (CNRS), Institut National de Recherche pour l’Agriculture, l’Alimentation et l’Environnement (INRAe) and Université de Tours.

## Author contributions

JLM, AKE and JAJB designed the experiments. BD performed stereotaxic surgeries for selective ablation experiments; BD, ADG, JLM, and JAJB performed behavioral testing. JG performed qRT-PCR experiments. ADG performed stereotaxic surgeries, fiber implantations, opto-stimulation during behavior and fos immunochemistry for optogenetic experiments; MF quantified fos expression. BD, JG, MF, ADG, JLM, AKE and JAJB analyzed the data. JLM, AKE, JAJB supervised the project and raised funding. JLM, AKE and JAB wrote the article. All authors discussed the findings, edited and contributed to the final version of the manuscript.

## Competing interests

The authors declare no conflict of interest.

## Supplementary methods

### Genotyping

Mice generated by the crossing of *Drd1a-Cre^+/-^* mice with iDTR^+/+^ mice were genotyped for the presence of the two transgenes using PCR. Drd1a-Cre: forward 5’ GCC-TGG-AGT-GAG-AAC-GAT-GTA-TCT-T 3’ and reverse 5’ TTT-TGG-TGT-ACG-GTC-AGT-AAA-TTG-G 3’. iDTR: forward 5’AAA-GTC-GCT-CTG-AGT-TGT-TAT 3’, reverse for mutated allele 5’ GCG-AAG-AGT-TTG-TCC-TCA-ACC 3’ and reverse for WT allele 5’ GGA-GCG-GGA-GAA-ATG-GAT-ATG 3’.

### Behavioral experiments

Experiments were performed in independent, dedicated cohorts of naïve mice. Behavioral assays in SPN-ablated mice were performed successively in the same cohort of mice, and testing order was chosen to minimize the incidence of anxiety. In experiments exploring the effects of VU0155041 treatment in NAc D1R-SPN mice, social interaction, stereotypies and 3-chamber test were also performed successively (see timeline in Figure 4B). In optogenetic experiments, social interaction was assessed on days 1,4,7,14 and 21 (see timeline in Figure 5B). Experiments were conducted and analyzed blind to genotype and experimental condition.

### Social behaviors

#### Direct social interaction test

The experimental protocol was adapted from (1, 2). As previously described (3–5), on testing day, the experimental mouse and an unfamiliar congener (wild-type C57BL/6 interactor mouse, age- and sex-matched) were introduced in one of 4 square arenas (40 x 40 cm, separated by 40 cm-high opaque grey Plexiglas walls) for 10 min (15 lx). The total amount of time spent in nose contact (nose-to-nose, nose-to-body or nose-to-anogenital region), the number of these contacts, the time spent in paw contact and the number of these contacts, grooming episodes (allogrooming), notably ones occurring immediately (<5s) after a social contact, as well as the number of following episodes were scored *a posteriori* on video recordings (EthoVision® XT, Noldus, Wageningen, the Netherlands) using an ethological keyboard (Labwatcher®, View Point, Lyon, France) by trained experimenters and individually for each animal. The mean duration of nose and paw contacts was calculated from these data (1, 2, 6). When social interaction was tested several times, each experimental mouse met a different interactor mouse for each test.

#### Three-chamber social preference test

The experimental protocol was adapted from (7). As previously described (3–5), the test apparatus consisted of a transparent acrylic box (exterior walls blinded with black plastic film); partitions divided the box into three equal chambers (40 x 20 x 22.5 cm). Two sliding doors (8 x 5 cm) allowed transitions between chambers. Cylindrical wire cages (18 x 9 cm, 0.5 cm diameter-rods spaced 1 cm apart) were used to contain the mouse interactor and object (soft-toy mouse). The test was performed in low-light conditions (15 lx) to minor anxiety. Stimulus wild-type mice were habituated to confinement in wire cages for 2 days before the test (20 min/day). On testing day, the experimental animal was introduced to the middle chamber and allowed to explore the whole apparatus for a 10-min habituation phase (wire cages empty) after the sliding doors were raised. The experimental mouse was then confined back in the middle-chamber while the experimenter introduced an unfamiliar wild type age and sex-matched animal into a wire cage in one of the side-chambers and a soft toy mouse (8 x 10 cm) in the second wire cage as a control for novelty. Then the experimental mouse was allowed to explore the apparatus for a 10-min interaction phase. The time spent in each chamber, the time spent in nose contact with each wire cage (empty: habituation; containing a mouse or a toy: interaction), as well as the number of these nose contacts were scored *a posteriori* on video recordings (EthoVision® XT, Noldus, Wageningen, the Netherlands) using an ethological keyboard (Labwatcher®, View Point, Lyon, France) by trained experimenters. The mean duration of nose contacts was calculated from these data (1, 2, 6). Preference ratio was calculated as follows: Time in nose contact with the mouse / (Time in nose contact with the mouse + Time in nose contact with the object) x 100. The relative position of stimulus mice (versus toy) was counterbalanced between groups.

### Stereotyped behaviors

#### Motor stereotypies

The experimental protocol was adapted from (7, 8). To detect spontaneous motor stereotypies in mutant versus wild-type animals, mice were individually placed in clear standard home cages (21×11×17 cm) filled with 3-cm deep fresh sawdust for 10 min, as described previously (7, 9). Light intensity was set at 30 lux. Trained experimenters scored numbers of head shakes, as well as rearing, burying, grooming, circling episodes and total time spent burying by direct observation.

#### Marble-burying

Marble burying was used as a measure of perseverative behavior (7, 8). Mice were introduced individually in transparent cages (21×11×17 cm) containing 20 glass marbles (diameter: 1.5 cm) evenly spaced on 4-cm deep fresh sawdust. To prevent escapes, each cage was covered with a filtering lid. Light intensity in the room was set at 40 lux. The animals were removed from the cages after 15 min, and the number of marbles buried more than half in sawdust was quoted.

#### Y-maze exploration

Spontaneous alternation behavior was used to assess perseverative behavior (10, 11). Each Y-maze (Imetronic, Pessac, France) consisted of three connected Plexiglas arms (15×15×17 cm) covered with distinct wall patterns (15 lx). Floors were covered with lightly sprayed fresh sawdust to limit anxiety. Each mouse was placed at the center of a maze and allowed to freely explore this environment for 5 min. The pattern of entries into each arm was quoted on video-recordings (EthoVision® XT, Noldus, Wageningen, the Netherlands). Spontaneous alternations (SPA), i.e. successive entries into each arm forming overlapping triplet sets, alternate arm returns (AAR) and same arm returns (SAR) were scored, and the percentage of SPA, AAR and SAR was calculated as following: total / (total arm entries −2) * 100.

### Motor function

#### Nest building

The protocol was adapted from (12, 13). As previously described (3, 14), experimental mice were single-housed overnight (16 hrs) in a clear standard cage (21 × 11 × 17 cm) provided with a single housing-device (red plastic igloo; SDS Mazuri, Argenteuil, France) with three openings. A block of nesting material was placed in the opposite end of the cage. Each cage was scored by adding the number of openings covered (1, 2 or 3) with nesting material to the condition of this material: 1 for initiation of shredding and 2 for a totally shredded nesting block.

#### Accelerating rotarod task

The apparatus was a rotarod (Ugo Basile, Gemonio, Italy) accelerating from 4 to 40 rpm in 5 min (40 lx). The experimental protocol was described previously (15, 16). On day 1, mice were habituated to rotation on the rod at 4 rpm, until they were able to stay more than 180 s. From day 2 to day 5, mice were tested for three daily trials (60 s intertrial). Each trial started by placing the mouse on the rod and beginning rotation at constant 4 rpm-speed for 60 s. Then the accelerating program was launched, and trial ended for a particular mouse when falling off the rod. Time stayed on the rod was automatically recorded.

### Anxiety-like behavior

#### Novelty-suppressed feeding

The protocol was adapted from (17). As described previously (3–5), novelty-suppressed feeding (NSF) was measured in 16-hr food-deprived mice, isolated in a standard housing cage for 30 min before individual testing. Three pellets of ordinary lab chow were placed on a white tissue in the center of one of 4 square arenas (50 x 50 cm, separated by 35cm-high opaque grey Plexiglas walls), filled with a 15-cm layer of sawdust and lit at 60 lx. Each mouse was placed in a corner of an arena and allowed to explore for a maximum of 15 min. Latency to feed was measured as the time necessary to bite a food pellet. Immediately after an eating event, the mouse was transferred back to home cage (free from cage-mates) and allowed to feed on lab chow for 5 min. The amount of food consumed in the home cage was measured.

#### Elevated Plus Maze

The protocol was described previously (15, 16). The EPM was a plus-shaped maze elevated 42 cm from base, with black Plexiglas floor, consisting of two open and two closed arms (31 x 7 cm each) connected by a central platform (7 x 7 cm). The walls of the closed arms were made of 17 cm-high opaque gray plexiglas.

Light intensity in open arms was set at 15 lx. The apparatus was placed over an infrared-lit platform. The movement and location of the mice were analyzed by an automated tracking system (EthoVision® XT, Noldus, Wageningen, the Netherlands). All sessions were videotaped for further analyses. Experimental mice were exposed to the EPM for 5 min at 15 lx. Anxiety-like behavior was assessed by spatiotemporal and ethological measures (15, 18). Automatically assessed spatiotemporal parameters included the distance traveled, time spent and number of entries in closed and open arms, and related distance, time and activity ratios (distance or time spent or number of entries in open arms/total distance or time spent or number of entries in arms). The time spent in the distal part of the open arms was measured to evaluate risk assessment behavior. Ethological measures were scored manually on video recordings and included the frequency of stretched attend postures (SAP; exploratory posture in which the body is stretched forward but the animal’s hind paws remain in position, followed by retraction to original position), flat back approaches (FBA; number of slow forward explorations with the body stretched) and head dips (HD; an exploratory behavior in which the animal scans over the sides of the maze towards the floor).

### Real-time quantitative PCR analysis

Real-time quantitative Polymerase Chain Reaction (qRT-PCR) analysis was performed on brain samples as described previously (4, 5). Brains were removed and placed into a brain matrix (ASI Instruments, Warren, MI, USA). Nucleus accumbens (NAc), medial caudate putamen (mCPu), ventral pallidum/olfactory tubercle (VP/Tu), and ventral tegmental area/substancia nigra pars compacta (VTA/SNc) were punched out/dissected from 1mm-thick slices (see Fig. S1). Tissues were immediately frozen on dry ice and kept at −80°C until use. For each structure of interest, genotype and condition, samples were processed individually (n=8). RNA was extracted and purified using the Direct-Zol RNA MiniPrep kit (Zymo research, Irvine, USA). cDNA was synthetized using the ProtoScript II Reverse Transcriptase kit (New England BioLabs, Évry-Courcouronnes, France). qRT-PCR was performed in quadruplets on a CFX384 Touch Real-Time PCR Detection System (Biorad, Marnes-la-Coquette, France) using iQ-SYBR Green supermix (Bio-Rad) kit with 0.25 µl cDNA in a 12 µl final volume in Hard-Shell Thin-Wall 384-Well Skirted PCR Plates (Bio-rad). Gene-specific primers were designed using Primer3 software to obtain a 100- to 150-bp product; sequences are displayed in Table S1. Relative expression ratios were normalized to the level of actin and the 2^−ΔΔCt^ method was applied to evaluate differential expression level. Gene expression values differing of the mean by more than two standard deviations were considered as outliers and excluded from further calculations.

### Fos immunohistochemistry and cell counting

Sixty minutes after social interaction, mice were deeply anesthetized with avertin (2,2,2-tribromoethanol, Sigma-Aldrich T48402, 2-methyl-2-butanol, Sigma-Aldrich 240486) and transcardially perfused with PBS 0.01M followed by 4% paraformaldehyde. Brains were removed and post-fixed in 4% paraformaldehyde. Cryoprotection was performed in 30% sucrose and brains were frozen and kept at − 80°C. Coronal sections (30μm) were performed on cryostat (Leica CM3050S) and kept in cryoprotective solution at −20°C (30% glycerol, 30% ethylene glycol, 40% PBS 0.01M pH 7.4).

Sections were incubated 1h with blocking solution (10% normal horse serum, 0.1% Triton X100, 0.1% sodium azide, PBS 0.01M pH 7.4) and overnight at +4°C with rabbit anti-Fos (1:1500, Cell Signaling Technology Cat# 2250, RRID:AB_2247211) and chicken anti-GFP (1:2000, Abcam Cat# ab13970, RRID:AB_300798). Appropriate secondary fluorescent antibodies were applied over 3h at room temperature (1:200, donkey anti-rabbit Alexa-647 Jackson Immunoresearch 711-605-152, RRID: AB_2492288; 1:400, goat anti-chicken Alexa-488 Invitrogen A-11039, RRID: AB_2534096). Finally, all sections were stained with Hoechst 33342 (1:500, Life Technologies H3570) and mounted (Fluorsave™ Millipore 345789).

Images were collected by epifluorescence microscopy (Zeiss Axio Zoom.V16, Hamamatsu ORCA-Flash4.0). For each brain region, counting was performed in two non-adjacent sections with ImageJ 2.3.0 software. Cell counts were normalized to the area of each brain region.

## Supplementary Figures

**Figure S1.**
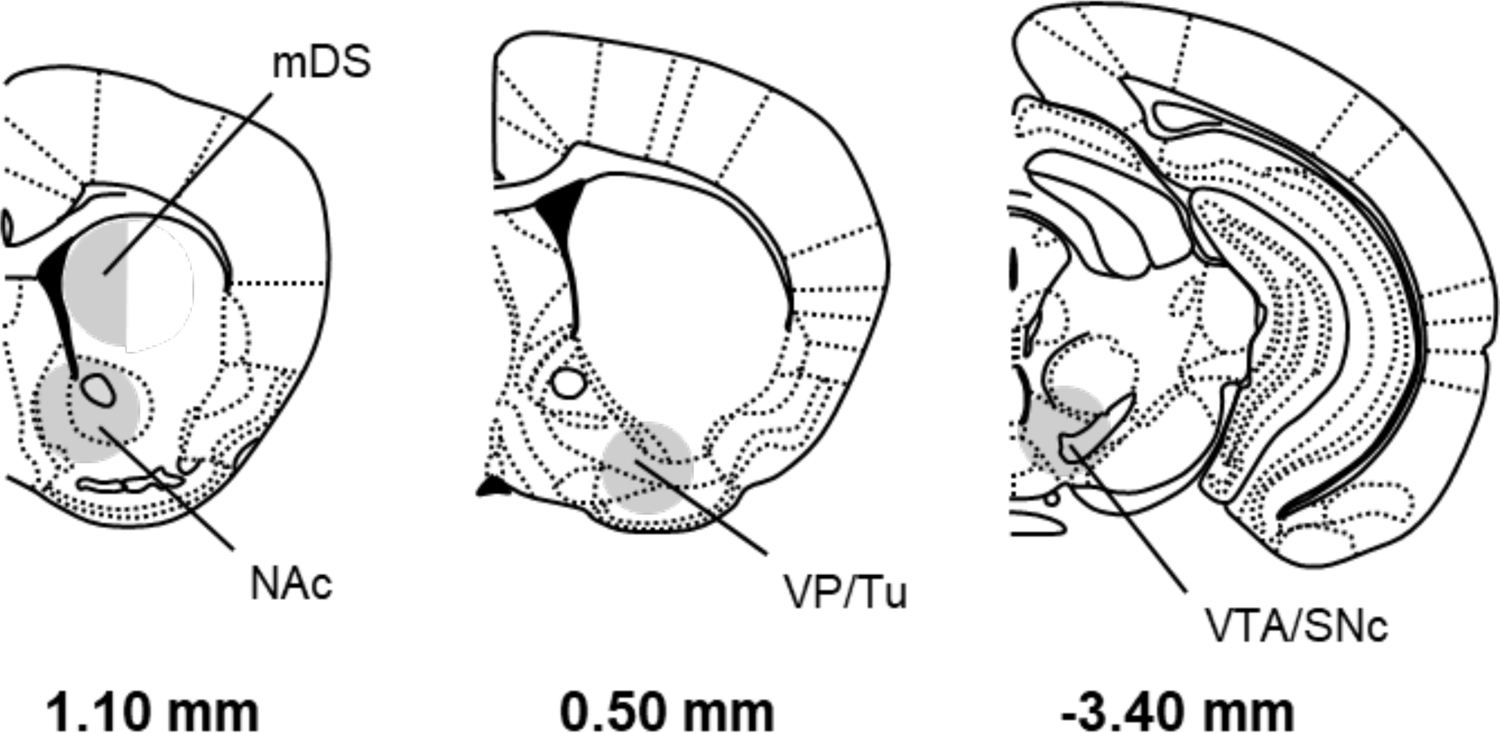
Schematic representations depict brain regions dissected for gene expression study and western blot experiments. NAc, mCPu, VP/Tu and VTA/SNc were punched on 1-mm thick brain slices (CPu: one punch/side, ᴓ 2 mm, medial half only; NAc, VP/Tu and VTA/SNc: one punch/side, ᴓ 1.25 mm). Coordinates refer to bregma. NAc: Nucleus accumbens; mDS: medial dorsal striatum; VP/Tu: ventral pallidum/Olfactory tubercle; VTA/SNc: Ventral tegmental area/ Substancia nigra, pars compacta.

**Figure S2.**
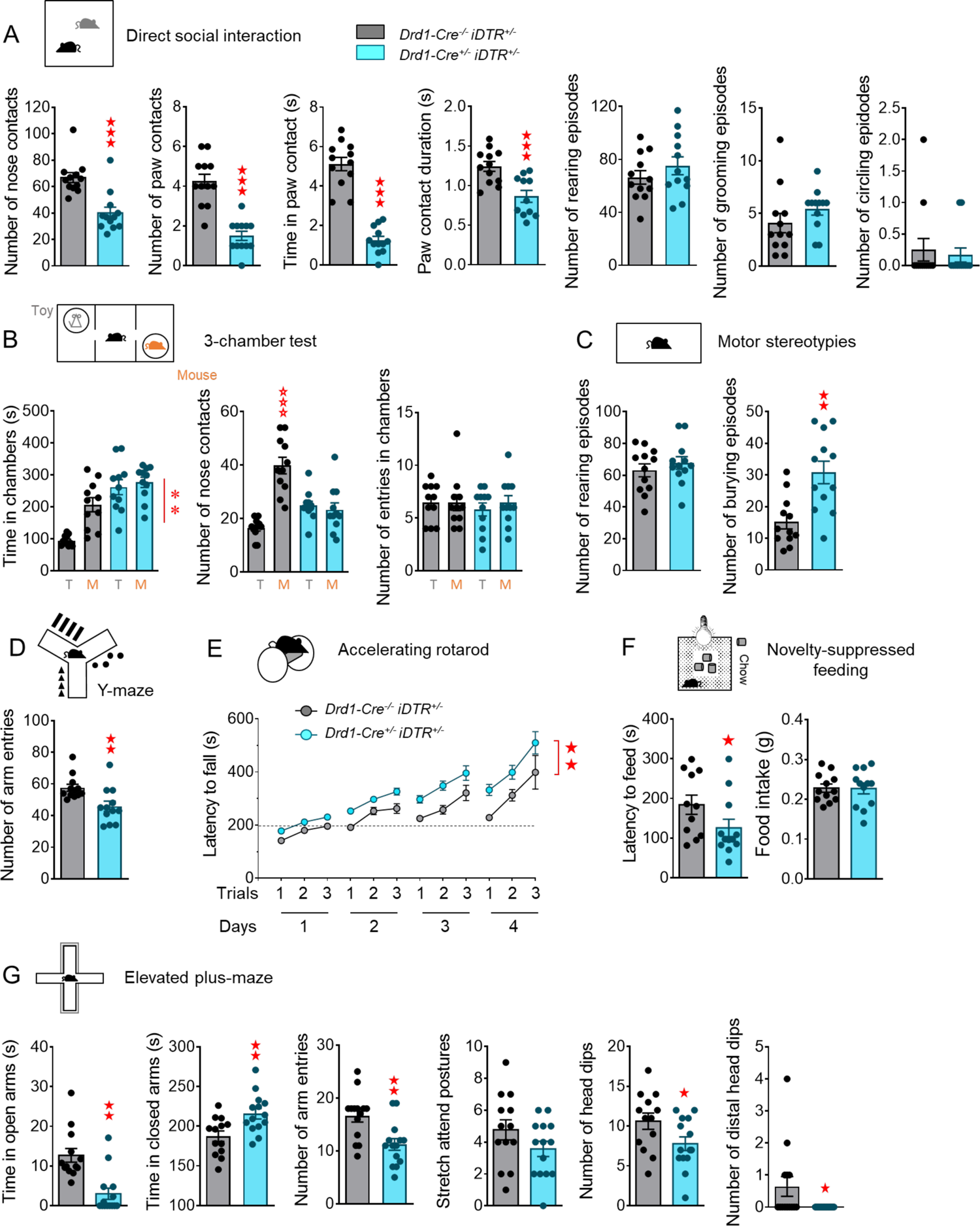
Behavioral consequences of ablating D1R-SPNs in the nucleus accumbens. See timeline of experiments and animal numbers in Figure 1A. **(A)** In the direct social interaction test, mice bearing D1R-SPN ablation (*D1-Cre^+/-^ iDTR^+/-^*) in the NAc displayed less frequent nose (*Ablation: F_1,22_=21.5, p<0.001)* and paw (*F_1,22_=42.9, p<0.0001)* contacts, reduced time spent in paw contact (*F_1,22_=98.5, p<0.0001)* and shorter paw contacts (*F_1,22_=15.1, p<0.001)*; their number of rearing episodes (used as an index of activity) was unchanged compared to controls (*D1-Cre^+/-^ iDTR^+/+^*), as were their numbers of grooming and circling episodes. **(B)** In the 3-chamber test, mice stayed longer in the chamber with the mouse (*Stimulus: F_1,20_=10.0, p<0.01)*; NAc D1R-SPN ablated mice failed to make more nose contacts with the mouse over the toy (*Stimulus x Ablation: F_1,20_=25.1, p<0.0001)*; their activity, as monitored by the number of entries in chambers, was similar to controls’. **(C)** During motor stereotypy test, NAc D1R-SPN deleted mice displayed similar number of rearing episodes (used as an index of activity) as sham controls, and increased number of burying episodes (*Ablation: F_1,22_=13.7, p<0.01)*. **(D)** During exploration of the Y-maze, mice with D1R-SPN deletion in the NAc showed decreased number of entries in arms (*Ablation: F_1,22_=9.0, p<0.01)*. **(E)** During the accelerated rotarod task, NAc D1R-SPN deleted mice performed better than *D1-Cre^+/-^ iDTR^+/+^* controls over sessions, as shown by increased latency to fall *(Session x Ablation: F_3,120_=8.2, p<0.001)*. **(F)** In the novelty-suppressed feeding test, lesioned mice took less time to eat in the middle of the arena *(Ablation: F_1,22_=4.4, p<0.05)* and ate similar amounts of food as controls. **(G)** In the elevated plus-maze, ablated mice spent less time in the open arms *(F_1,26_=10.0, p<0.01)* but increased time in closed arms *(F_1,26_=10.3, p<0.01)*, globally entered less in arms *(F_1,26_=11.2, p<0.01)*, displayed similar numbers of stretch attend postures and head-dipped less (*F_1,26_=5.9, p<0.05)*, notably at distal parts of the open arms (*F_1,26_=4.4, p<0.05)*, than sham controls. Results are shown as scatter plots and mean ± sem. Solid stars: ablation effect (one/three-way or Kruskal-Wallis ANOVA), asterisk: stimulus effect (two-way ANOVA), open stars: ablation x stimulus interaction (mouse versus object comparison). One symbol: p<0.05, two symbols: p<0.01; three symbols: p<0.001.

**Figure S3.**
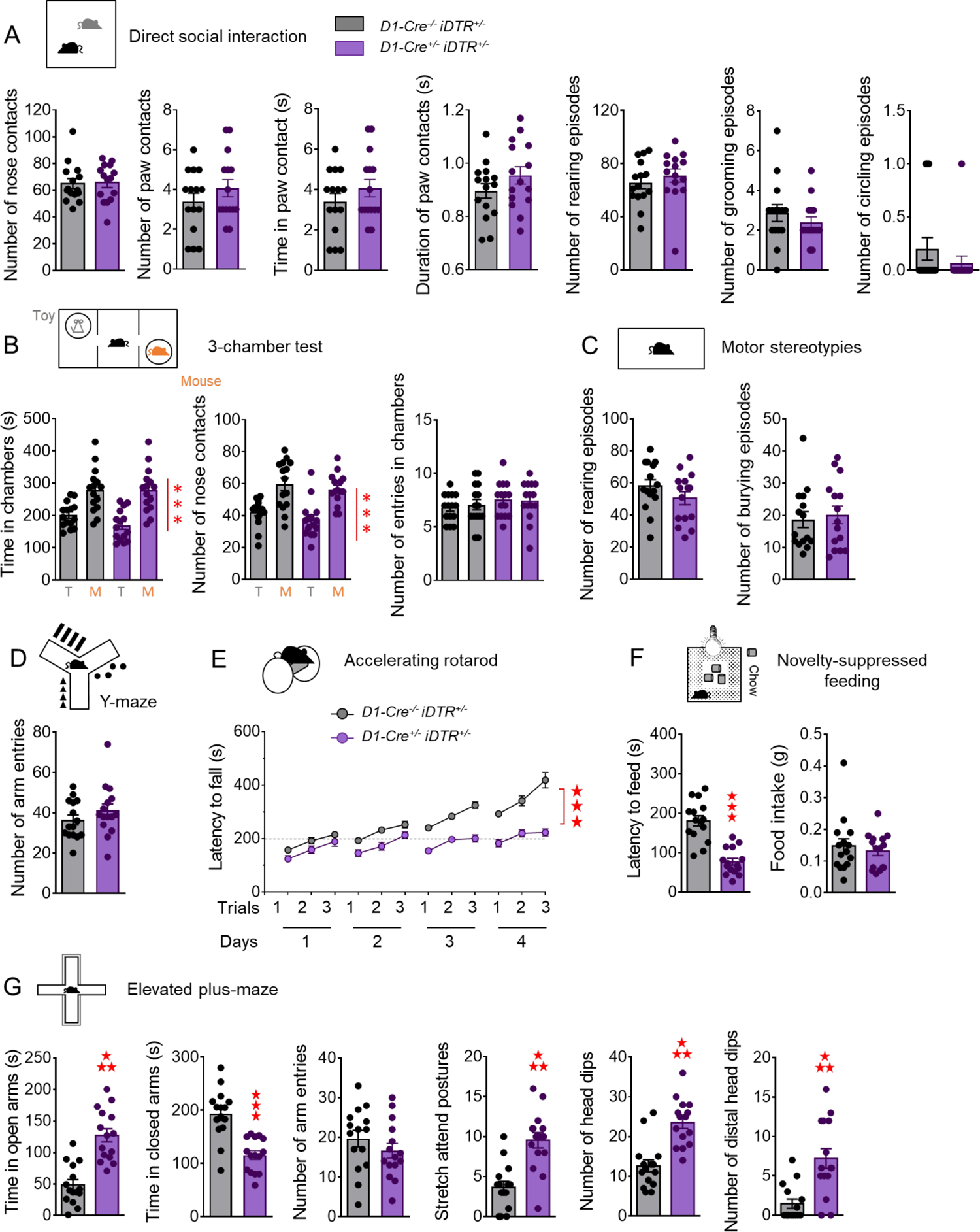
Behavioral consequences of ablating D1R-SPNs in the medial dorsal striatum. See timeline of experiments and animal numbers in Figure 2A. **(A)** In the direct social interaction test, mice bearing D1R-SPN ablation (*D1-Cre^+/-^ iDTR^+/-^*) in the mDS displayed similar number of nose and paw contacts, time spent in paw contact and mean duration of paw contacts, number of rearing (activity index), grooming and circling episodes compared to sham controls (*D1-Cre^+/-^ iDTR^+/+^*). **(B)** In the 3-chamber test, all the mice stayed longer in the chamber with the mouse (*Stimulus: F_1,28_=38.1, p<0.0001)* and made more frequent nose contacts with the mouse over the toy (*Stimulus: F_1,28_=30.4, p<0.0001)*; the activity of lesioned mice, as monitored by the number of entries in chambers, was similar to controls’. **(C)** When tested for motor stereotypies, mDS D1R-SPN deleted mice displayed similar number of rearing (used as an index of activity) and burying episodes as sham controls. **(D)** During exploration of the Y-maze, mice with D1R-SPN deletion in the mDS showed no difference in their number of arm entries as compared to controls. **(E)** During the accelerated rotarod task, mDS D1R-SPN deleted mice showed severe impairment in motor skill learning, as illustrated by maintained short latency to fall over sessions and trials *(Session x Trials x Ablation: F_6,168_=8.2, p<0.001)*. **(F)** In the novelty-suppressed feeding test, lesioned mice took less time to eat in the middle of the arena *(Ablation: F_1,28_=43.8, p<0.0001)* and ate similar amounts of food as controls. **(G)** In the elevated plus-maze, ablated mice spent less time in the open arms *(F_1,28_=34.7, p<0.0001)* but increased time in closed arms *(F_1,28_=28.0, p<0.0001)*, displayed similar numbers of arm entries and stretch attend postures but head-dipped less (*F_1,28_=26.1, p<0.0001)*, notably at distal parts of the open arms (*F_1,28_=17.2, p<0.001)*, than sham controls. Results are shown as scatter plots and mean ± sem. Solid stars: ablation effect (one/three-way or Kruskal-Wallis ANOVA), asterisk: stimulus effect (two-way ANOVA). Three symbols: p<0.001.

**Figure S4.**
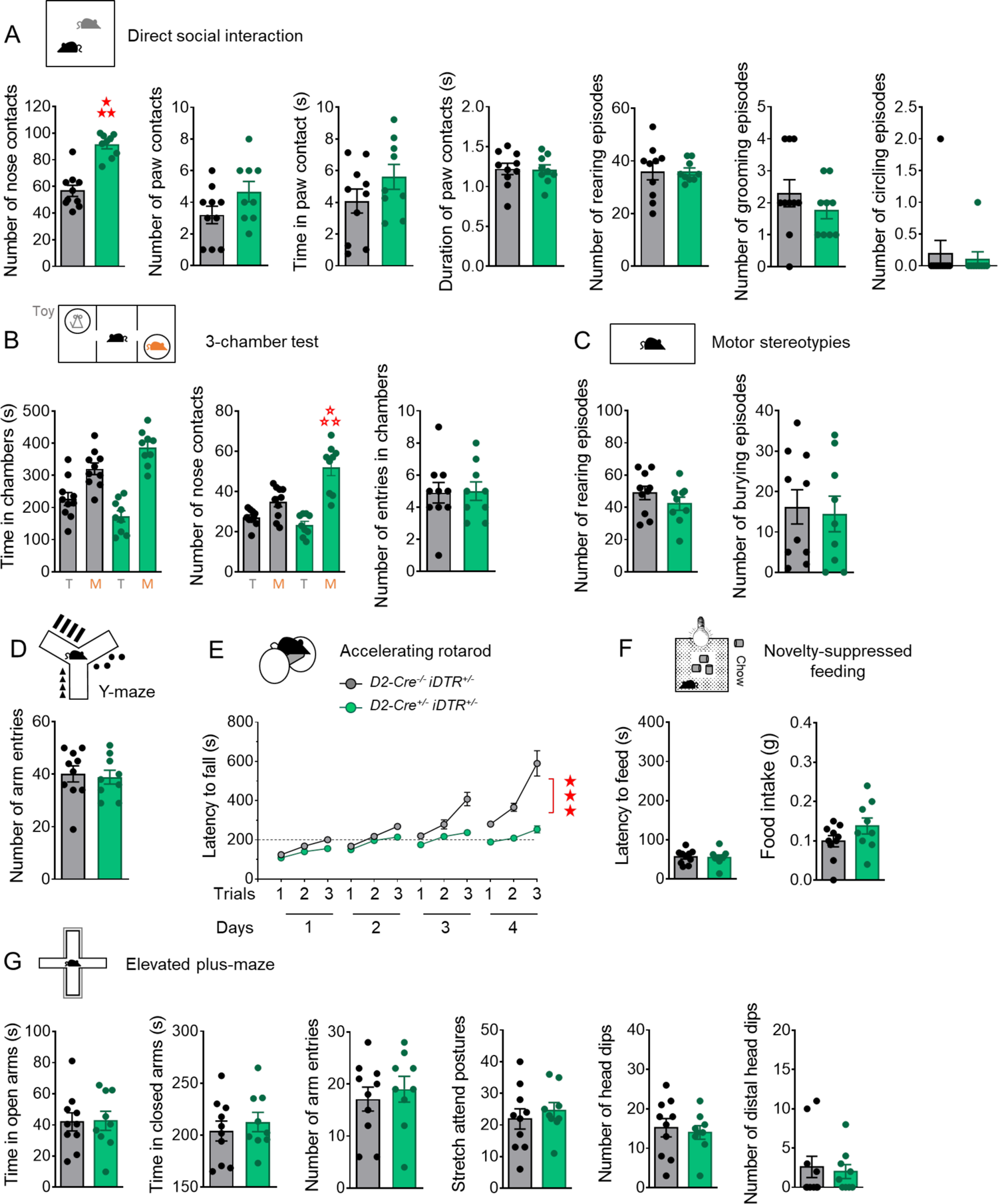
Behavioral consequences of ablating D2R-SPNs in the nucleus accumbens. See timeline of experiments and animal numbers in Figure 3A. **(A)** In the direct social interaction test, mice bearing D2R-SPN ablation (*D2-Cre^+/-^ iDTR^+/-^*) in the NAc displayed more frequent nose contacts (*Ablation: F_1,17_=74.8, p<0.0001*) and similar number, time spent in and duration of paw contacts; their number of rearing (an index of activity), grooming and circling episodes were also unchanged compared to sham controls (*D2-Cre^+/-^ iDTR^+/+^*). **(B)** In the 3-chamber test, all the mice stayed longer in the chamber with the mouse (*Stimulus: F_1,17_=5.5, p<0.05)*; ablated mice made significantly more frequent nose contacts with the mouse over the toy (*Stimulus x Ablation: F_1,17_=12.2, p<0.01*); the number of entries in chambers was similar between ablated and sham mice. **(C)** When tested for motor stereotypies, NAc D2R-SPN deleted mice displayed similar number of rearing (an index of activity) and burying episodes as sham controls. **(D)** During exploration of the Y-maze, mice with D2R-SPN deletion in the NAc showed made similar number of entries in arms as compared to controls. **(E)** During the accelerated rotarod task, NAc D2R-SPN deleted mice showed severe impairment in motor skill learning, as illustrated by maintained short latency to fall over sessions and trials *(Session x Trials x Ablation: F_6,102_=11.3, p<0.0001)*. **(F)** In the novelty-suppressed feeding test, lesioned mice took as long to eat in the middle of the arena and ate similar amounts of food than controls. **(G)** In the elevated plus-maze, NAc D2R-SPN ablated mice displayed similar time spent in open and closed arms, number of arm entries, stretch attend postures and head dips, including from the distal part of the maze, as sham controls. Results are shown as scatter plots and mean ± sem. Solid stars: ablation effect (one/three-way or Kruskal-Wallis ANOVA), open stars: ablation x stimulus interaction (two-way ANOVA, mouse versus object comparison). Three symbols: p<0.001.

**Figure S5.**
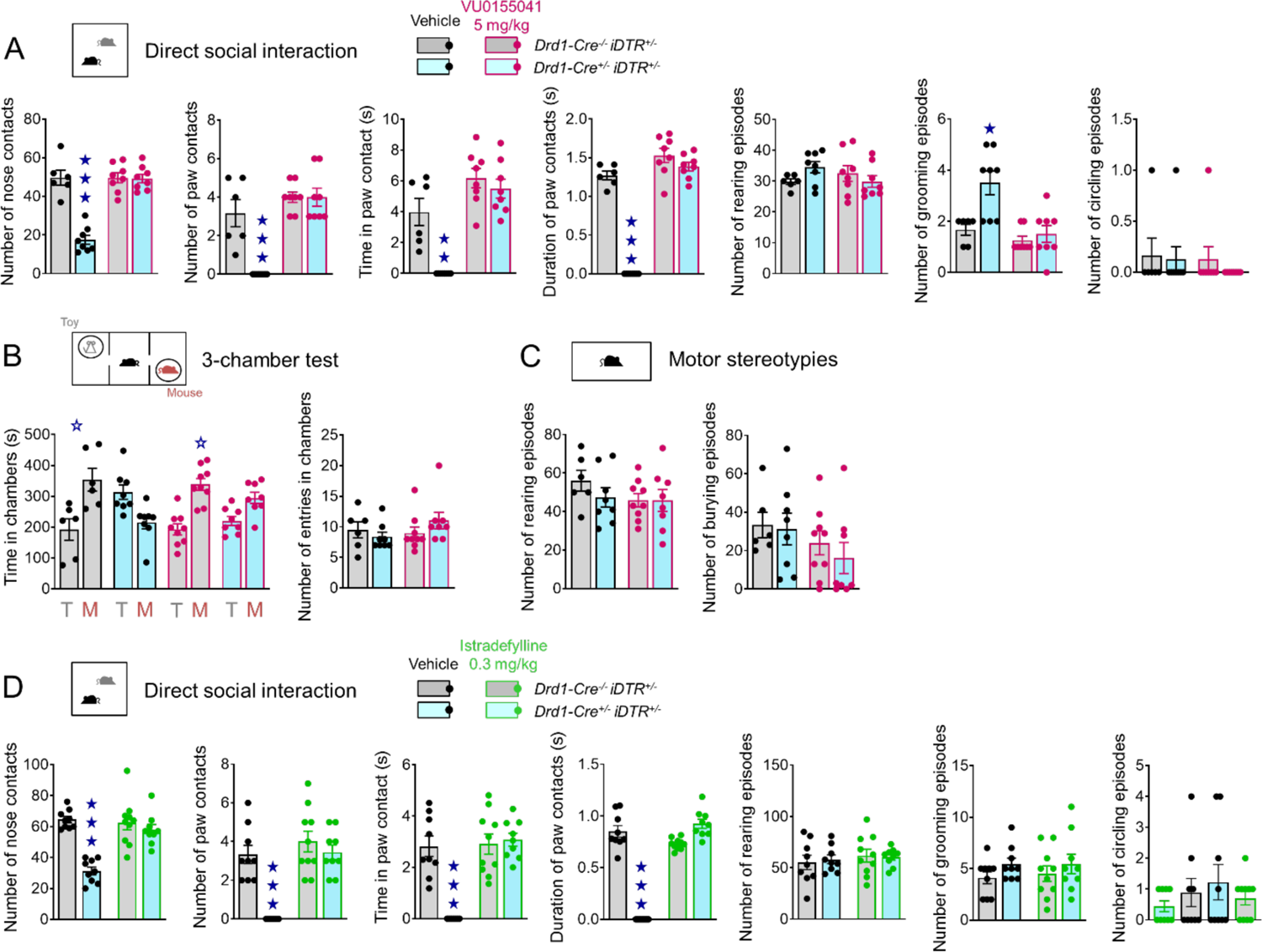
Behavioral consequences of D2R-SPN pharmacological inhibition in mice bearing D1R-SPN ablation in the nucleus accumbens. See timeline of experiments and animal numbers in Figure 4A and 4F. **(A)** In the direct social interaction test, chronic VU0155041 normalized the number of nose (*Treatment x Ablation: F_1,26_=34.8, p<0.0001)* and paw contacts (*T x A: F_1,26_=15.6, p<0.001)*, the time spent in paw contact (*T x A: F_1,26_=7.8, p<0.01)* and the duration of such contacts (*T x A: F_1,26_=83.9, p<0.0001)* of NAc D1R-SPN ablated mice; vertical activity and number of circling episodes were similar between ablated/ sham and VU0155041/vehicle-treated mice; chronic mGlu4 facilitation normalized the number of grooming episodes in lesioned mice (*T x A: F_1,26_=5.8, p<0.05)*. **(B)** In the 3-chamber test, only sham mice, saline or VU0155041-treated, spent more time in the chamber with the mouse (Stimulus x *Treatment x Ablation: F_1,27_=4.5, p<0.05);* the number of entries in chambers was similar between all groups. **(C)** During scoring of locomotor stereotypies, all mice displayed similar numbers of rearing and burying episodes. **(D)** In the direct social interaction test, acute administration of istradefillyne normalized the number of nose and paw contacts, the time spent in and duration of paw contacts of NAc D1R-SPN ablated mice; the number of rearing, grooming and circling episodes were similar between all groups in this experiment. Results are shown as scatter plots and mean ± sem. Solid stars: ablation x treatment interaction (two-way ANOVA), open stars: ablation x treatment x stimulus interaction (three-way ANOVA, mouse versus object comparison). One symbol: p<0.05, two symbols: p<0.01; three symbols: p<0.001.

**Figure S6.**
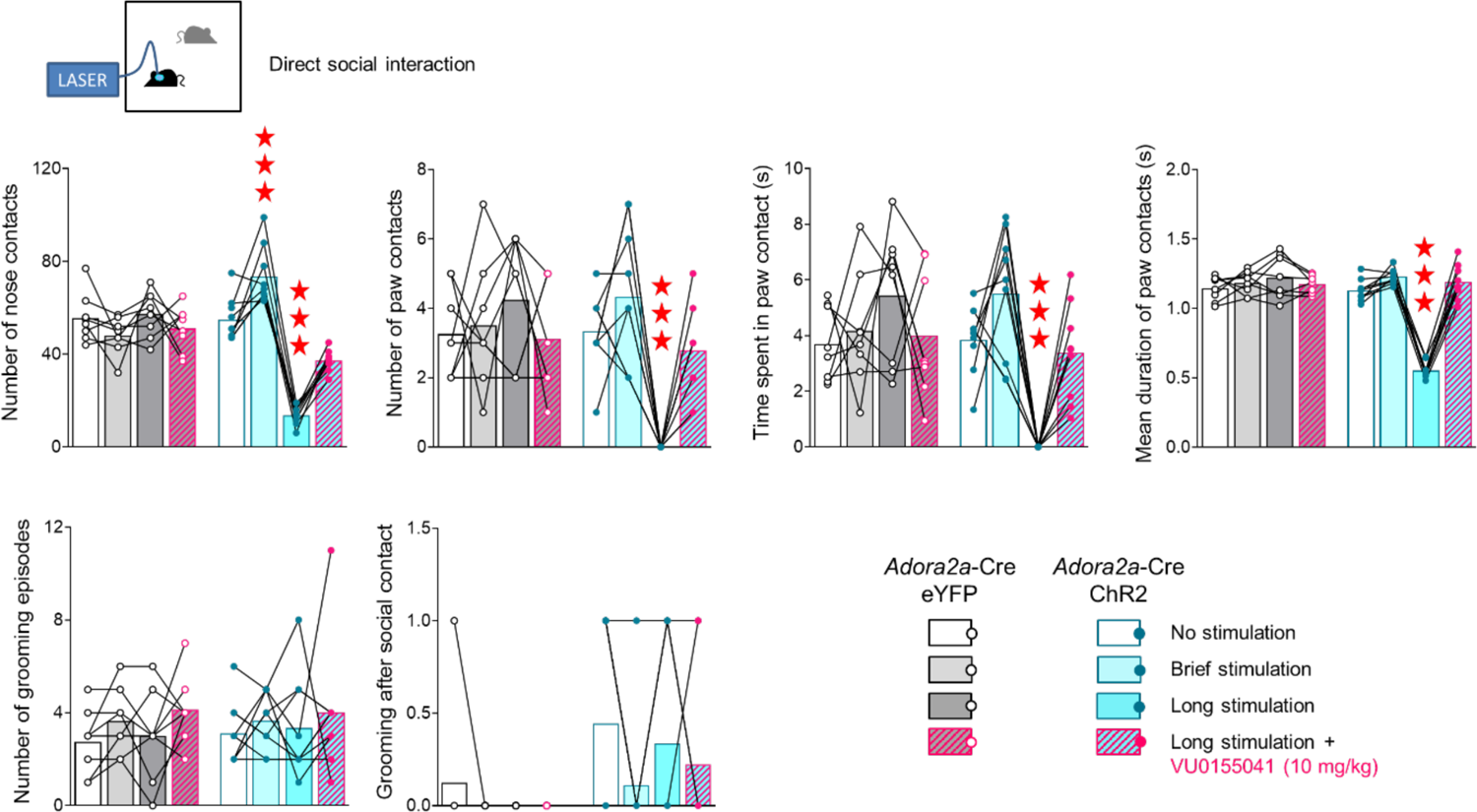
Effects of optical stimulation of D2R-SPN in the nucleus accumbens on social interaction. See timeline of experiments and animal numbers in Figure 5A. In the direct social interaction test, brief optical D2R-SPN stimulation in the NAc increased the number of nose contacts in Adora2a-cre-ChR2 mice (*Viral construct x Condition: F_3,45_=42.8, p<0.0001*). Prolonged stimulation reduced the number of nose (*V x C: F_3,45_=42.8, p<0.0001)* and paw contacts (*V x C: F_3,45_=8.9, p<0.0001)* as well as the time spent in paw contacts (*V x C: F_3,45_=9.9, p<0.0001)* and their duration (*V x C: F_3,45_=100.7, p<0.0001)*, with acute VU0155041 treatment preventing these deleterious effects. The number of grooming episodes, including those occurring immediately after a social contact, was not influenced by optogenetic manipulation nor pharmacological treatment. Results are shown as scatter plots and mean ± sem. Solid stars: optical stimulation effect (two-way ANOVA, compared to no light condition in *Adora2a-cre-*YFP mice). One symbol: p<0.05, two symbols: p<0.01; three symbols: p<0.001.

## Legends to supplementary tables

**Table S1.** List of primers used for qRT-PCR

**Table S2. Transcriptional consequences of ablating D1R-SPNs in the nucleus accumbens.** Data are expressed as fold change versus *Drd1-Cre^-/-^ iDTR^+/-^* control group (mean ± SEM). Two tailed t-tests were performed on transformed data (see Material and Methods). An adjusted p value was calculated using Benjamini-Hochberg correction for multiple testing. Significant regulations of gene expression are highlighted in bold and filled in red for significant up-regulation or in blue for significant down-regulation. NAc: nucleus accumbens, mDS: medial dorsal striatum, SN/VTA: substantia nigra/ventral tegmental area.

**Table S3. Transcriptional consequences of ablating D1R-SPNs in the medial dorsal striatum.** Data are expressed as fold change versus *Drd1-Cre^-/-^ iDTR^+/-^* control group (mean ± SEM). Two tailed t-tests were performed on transformed data (see Material and Methods). An adjusted p value was calculated using Benjamini-Hochberg correction for multiple testing. Significant regulations of gene expression are highlighted in bold and filled in red for significant up-regulation or in blue for significant down-regulation. NAc: nucleus accumbens, mDS: medial dorsal striatum, SN/VTA: substantia nigra/ventral tegmental area.

**Table S4. Transcriptional consequences of ablating D2R-SPNs in the nucleus accumbens.** Data are expressed as fold change versus *Drd2-Cre^-/-^ iDTR^+/-^* control group (mean ± SEM). Two tailed t-tests were performed on transformed data (see Material and Methods). An adjusted p value was calculated using Benjamini-Hochberg correction for multiple testing. Significant regulations of gene expression are highlighted in bold and filled in red for significant up-regulation or in blue for significant down-regulation. NAc: nucleus accumbens, mDS: medial dorsal striatum, SN/VTA: substantia nigra/ventral tegmental area.

**Table S5.** Transcriptional consequences of D2R-SPN pharmacological inhibition in mice bearing D1R-SPN ablation in the nucleus accumbens. Data are expressed as fold change versus saline *Drd1-Cre^-/-^ iDTR^+/-^* group (mean ± SEM). Two tailed t-tests were performed on transformed data (see Material and Methods). An adjusted p value was calculated using Benjamini-Hochberg correction for multiple testing. Significant regulations of gene expression are highlighted in bold and filled in red for significant up-regulation or in blue for significant down-regulation. NAc: nucleus accumbens, mDS: medial dorsal striatum, SN/VTA: substantia nigra/ventral tegmental area.

## Notes

### Competing Interest Statement

The authors have declared no competing interest.

